# Placental neurosteroids shape cerebellar development and social behaviour

**DOI:** 10.1101/730150

**Authors:** Claire-Marie Vacher, Jiaqi J. O’Reilly, Jacquelyn Salzbank, Helene Lacaille, Dana Bakalar, Sonia Sebaoui-Illoul, Philippe Liere, Cheryl Clarkson-Paredes, Toru Sasaki, Aaron Sathyanesan, Yuka Imamura Kawasawa, Anastas Popratiloff, Kazue Hashimoto-Torii, Vittorio Gallo, Michael Schumacher, Anna A. Penn

## Abstract

Compromised placental function or premature loss has been linked to diverse neurodevelopmental disorders ^1,2^. The placenta is the first functional foetal endocrine organ, but the direct impact of placental hormone loss on foetal brain in late gestation has not been empirically tested. Allopregnanolone (ALLO) is a non-glucocorticoid, progesterone derivative that acts as a positive modulator of GABA-A receptor activity^3^ with the potential to alter critical GABA-mediated developmental processes ^4,5^. To directly test the role of placental ALLO, we generated a novel mouse model in which the gene encoding the synthetic enzyme for ALLO (*Akr1c14*) is specifically deleted in trophoblasts using a tissue-specific Cre-Lox strategy. ALLO concentrations are significantly decreased in late gestation in placenta and brain when placental *Akr1c14* is removed, indicating placenta as the primary gestational ALLO source. We now demonstrate that targeted placental ALLO loss leads to permanent changes in brain development in a sex- and regionally-specific manner. Placental ALLO insufficiency led to male-specific cerebellar white matter (WM) abnormalities characterized by excess myelination with increased myelin protein expression, similar to changes reported in boys with autism spectrum disorders (ASD)^6,7^. Behavioural testing of these mice revealed increased repetitive behaviour and sociability deficits, two hallmarks of ASD, only in male offspring with placental ALLO insufficiency. Notably, a strong positive correlation was seen between the cerebellar contents of myelin basic protein (MBP) and the severity of ASD-like behaviours. A single injection of ALLO during gestation was sufficient to rescue both cerebellar MBP levels and aberrant behaviours. This study reveals a new role for a placental hormone in shaping specific brain structures and behaviours, and suggests that identifying placental hormone insufficiency or preterm loss may offer novel therapeutic opportunities to prevent later neurobehavioural disorders.

## MAIN

During foetal development, *Akr1c14*, the gene encoding the ALLO synthesis enzyme (3α-HSD, 3α-hydroxysteroid dehydrogenase), is primarily expressed in the placenta, with up to 60 times higher expression than in the mouse brain (Fig. 1a,b), similar to the human maternal ALLO peak in the second half of gestation^8^. To assess the neurodevelopmental role of endogenous ALLO, we generated *Akr1c14-floxed* mice (Fig. 1c) and crossed them with Cre-transgenic mice in which Cre recombinase is driven by the human Cyp19a promoter, which in mouse is active only in placenta^9^. In the Akr1c14^Cyp19a^ knockout mice (abbreviated as plKO for placental conditional KO), *Akr1c14* transcript was significantly reduced in the placenta, but not brain (Fig. 1d; Extended Data Fig. 1). Direct steroid measurements revealed a ∼50% ALLO reduction in both placenta and brain compared to littermate controls (Fig. 1e), consistent with placental provision of ALLO to the foetal brain. ALLO precursors and other neuroactive steroids in the synthesis pathway were not altered (Extended Data Fig. 2).

**Figure 1.**
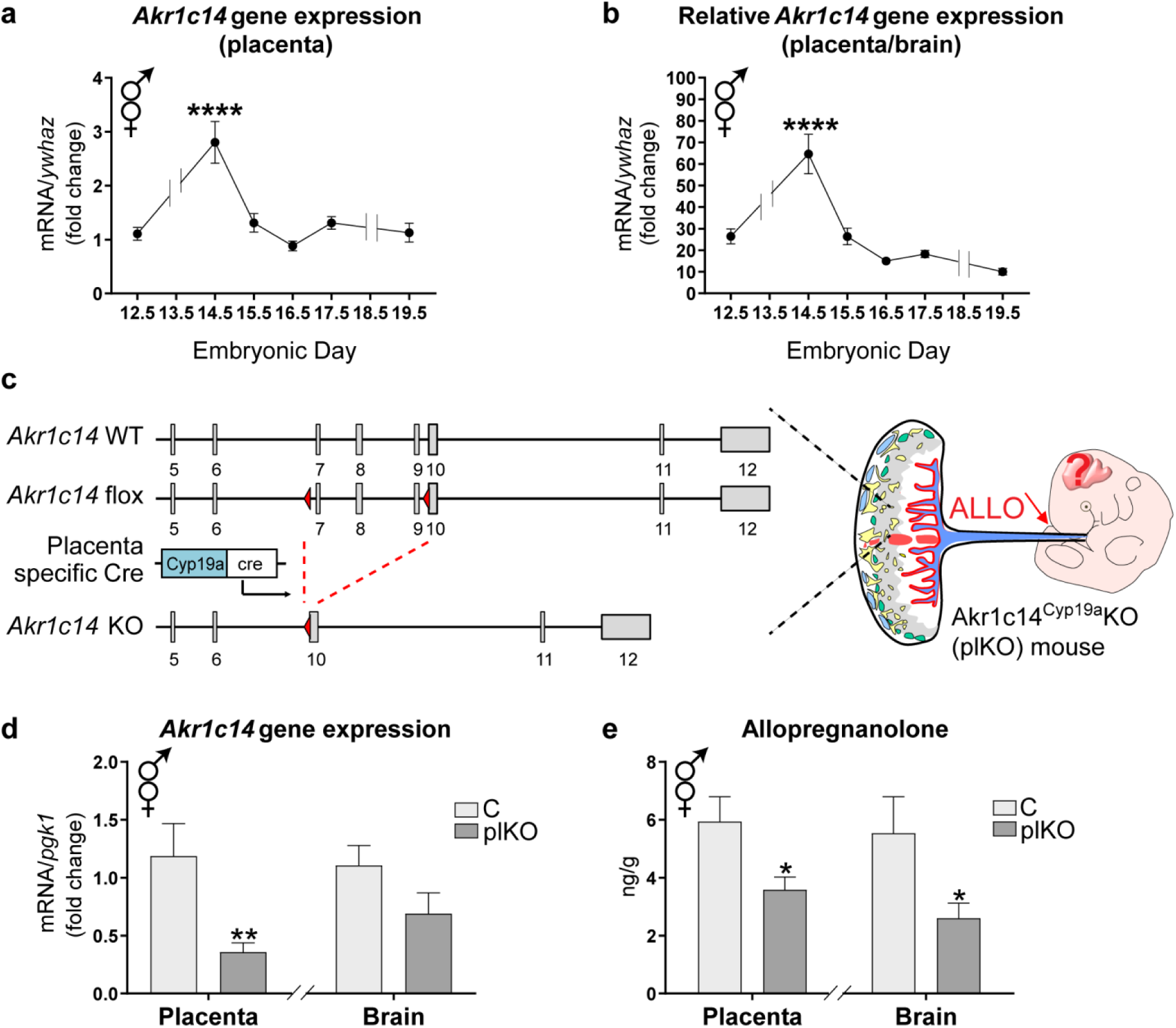
Conditional deletion of *Akr1c14* in Cyp19a-expressing trophoblasts results in reduced ALLO levels in the foetal brain. **a.** qRT-PCR for *Akr1c14* in wild-type (WT) mice. There was no significant difference between males and females. **b.** Relative *Akr1c14* gene expression in placenta vs brain in WT mice. Data presented as mean fold changes ±SEM (n=7-12 embryos/time point). One-way ANOVA with Dunnett’s multiple comparisons. ****p<0.0001 as compared with E12.5 value. *ywhaz*, Tyrosine 3-Monooxygenase/Tryptophan 5-Monooxygenase Activation Protein Zeta. **c.** *Akr1c14* genetic locus before and after recombination. Exons 7-9 are conditionally targeted. Cyp19a promoter drives expression of Cre in the placenta only. LoxP sites are indicated by red triangles, and; Exons 5-12 by grey boxes. **d.** qRT-PCR for *Akr1c14* normalized to *pgk1* at E17.5. Data presented as means ± SEM (n=7-9 samples/group). *Pgk1*, phosphoglycerate kinase 1. **e.** Mass spectrometry hormonal assays at E17.5. Data presented as means ± SEM (n=9-12 pools of 3 samples/group). *p<0.05, **p<0.01 (two-tailed unpaired Student’s t test with Welch’s correction).

Comparison of plKO mice to littermate controls (C) provides the opportunity to examine foetal and long-term neurodevelopmental effects of placental hormone insufficiency. To define the most prominent long-term molecular alterations associated with placental ALLO insufficiency in a non-biased way, we performed comparative RNA sequencing analysis in cerebral cortex, hippocampus and cerebellum of C and plKO mice at postnatal day (P) 30 in both sexes. Cutoff criteria used to identify significant differentially expressed genes (DEGs) were validated by RT-PCR and Western blots (Extended Data Fig. 3). Male cerebellum was the brain structure impacted most by placental ALLO loss as measured by the number of DEGs, which was 3 to 5× higher than in any other brain region (Extended Data Fig. 4a, b). Cerebellar DEGs were equally distributed in up- and down-regulated gene categories (Extended Data Fig. 4a-d). Ingenuity Pathway Analysis (IPA; Qiagen, Redwood City, CA) identified white matter (WM)-associated pathways as the top category of altered genes (p=0.0053 and 0.041 in males and females, respectively). Comparison of cerebellar DEGs to a previous published oligodendrocyte (OL) transcriptome^10^ revealed an overlap of 51 and 31 genes in males and females, respectively (Extended Data Fig. 4e,f). Striking sexual dimorphism was seen: male and female DEGs were qualitatively different and the majority of OL-related DEGs were up-regulated in plKO males but down-regulated in female plKO (Extended Data Fig. 4g,h).

Structural characterization of the cerebellar WM by electron microscopy in males (Extended Data Fig. 5a) revealed thicker myelin in plKO (g-ratio measurements, Fig. 2a,a’). In contrast, female plKO myelinated cerebellar fibers showed unchanged g-ratios (Extended Data Fig. 5b). Further investigation of myelin structural proteins revealed major sex-specific changes: in plKO males, MBP, MAG and MOG protein levels were all significantly increased (Fig. 2b,b’), while in plKO females, MBP cerebellar content was reduced (Extended Data Fig. 5c,c’), changes confirmed by RT-PCR and immunohistochemistry (Extended Data Fig. 5d,e).

**Figure 2.**
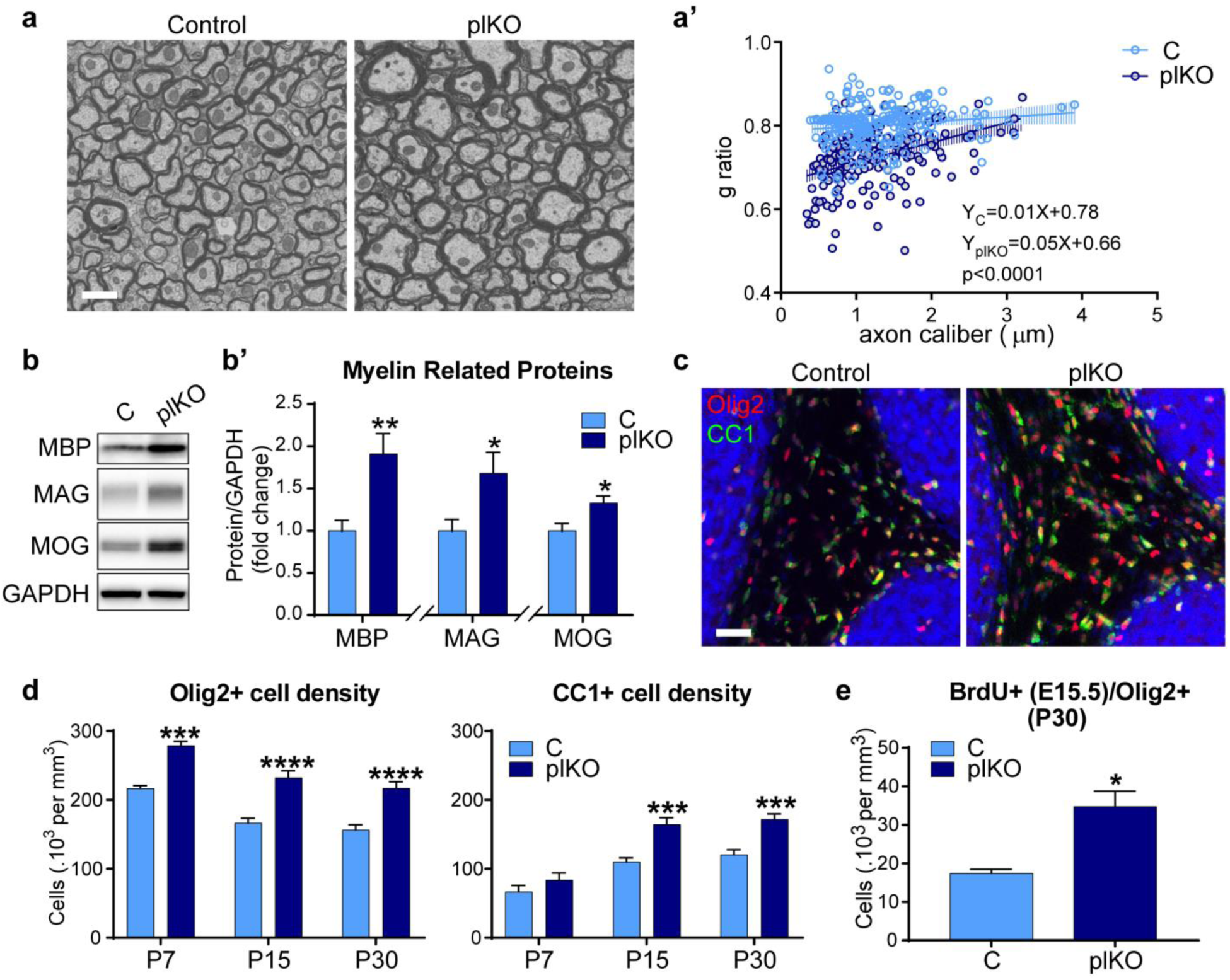
Placental ALLO insufficiency results in cerebellar WM impairment in males. **a.** Representative scanning electron microscopy acquisitions (20,000×) in inter-lobule VI-VII WM at P30. Scale bar, 3 μm. **a’.** Scatter plot of g ratios of >300 individual axons; data from one representative control and one representative plKO mouse. Fitted lines are linear regressions. **b-b’.** Western blot analysis of myelin related proteins in the cerebellum at P30. Data presented as mean fold changes ± SEM (n=6-28/group). *p<0.05; **p<0.005 (two-tailed unpaired Student’s t test with Welch’s correction). GAPDH, Glyceraldehyde 3-phosphate dehydrogenase; MAG, myelin associated glycoprotein; MBP, myelin basic protein; MOG, myelin oligodendrocyte glycoprotein. **c.** Immunofluorescent staining of Olig2 and CC1 in inter-lobule VI-VII WM. Scale bar, 100 μm. **d.** Quantification of Olig2+ and CC1+ cell density within the cerebellar WM. Data presented as means ± SEM. ***p<0.005, ****p<0.0001 (two-way ANOVA with Sidak’s multiple comparisons test). **e.** Quantification of BrdU+/Olig2+ cell density in the cerebellar WM at P30 (n=5/group). Data presented as means ± SEM. *p<0.05 (two-tailed unpaired Student’s t test with Welch’s correction).

Cerebellar myelination is a postnatal process^11^ so placental endocrine disruption leading to hypermyelination was unanticipated. However, the adjustment of myelin sheath thickness relies on the local density of myelin-competent oligodendrocytes (OLs) that are primarily produced before birth as OL progenitor cells (OPCs)^11^, at a time when ALLO is highest. ALLO is a potent positive allosteric modulator of GABA-A receptor and GABA-A signalling is associated with blockade of OPC proliferation^12,13^, so we tested the hypothesis that prenatal ALLO insufficiency enhances OLs production in males. We found that the density of OL lineage cells (Olig2^+^) and particularly that of mature OLs (CC1^+^) were significantly increased in the male plKO cerebellum at P30 compared to controls (Fig. 2c,d). Increased OPC proliferation during foetal life is suggested by the results of co-labelling with BrdU at E15.5 (marking cells dividing between E15.5-16.5) and Olig2 expression at P30. Additionally, within the OL lineage, a transient acceleration of the maturation from PDGFRα^+^ OPCs to mature CC1^+^ OLs was observed in the plKO mice at P15, when cerebellar myelination is starting (Extended Data Fig. 5f). Overall, these results are consistent with the ultrastructural analysis of myelin, and indicate immediate and long-term modifications of cerebellar OL lineage progression and OL maturation when placental ALLO is low. No volumetric change in the different cerebellar layers nor in Purkinje cell linear density was observed in the plKO mice (Extended Data Fig. 6).

Given the cerebellar WM alterations seen in plKO mice, we investigated the functional impact of these changes using standard neurobehavioural and cerebellum-specific tests. Cerebellar circuits provide precise spatiotemporal control of sensorimotor behaviour. Cerebellar contribution to cognitive functions has more recently been recognized ^14^. Thus, developmental disruption of cerebellar circuits either through neuronal or glial alterations may alter motor and cognitive performance. Mice were tested using the Erasmus ladder, an advanced, fully automated system for comprehensive real-time monitoring of cerebellar-mediated motor learning and behaviour ^15^. At baseline, male plKO mice exhibited a slightly altered walking pattern, with a longer stride length (Extended Data Fig. 7a,b), but their locomotion speed was no different than controls on the Erasmus ladder or in open field testing (Extended Data Fig. 7c,d). An associative cerebellar learning task pairing a tone with a ladder obstacle revealed no cerebellar motor learning deficit in male plKO mice (Extended Data Fig. 7e-h). Female plKOs exhibited no locomotor alterations (Extended Data Fig. 8). To further evaluate balance, motor coordination and learning, mice were tested on an accelerating rotarod. Male, but not female, plKO mice displayed longer latency to fall and increased terminal speed (Fig. 3a; Extended Data Fig. 9a,c,d). The learning rate of individual mice in the accelerating rotarod, given by a linear regression analysis, was also higher in male plKO mice (Extended Data Fig. 9a). The inclined beam walking test corroborated the enhanced locomotor and coordination performances seen in male plKO mice (Extended Data Fig. 9b). Similar locomotor enhancement, rather than impairment, has been described in mouse models of ASD and has been attributed to increased repetitive/stereotypical behaviour^16^.

**Figure 3.**
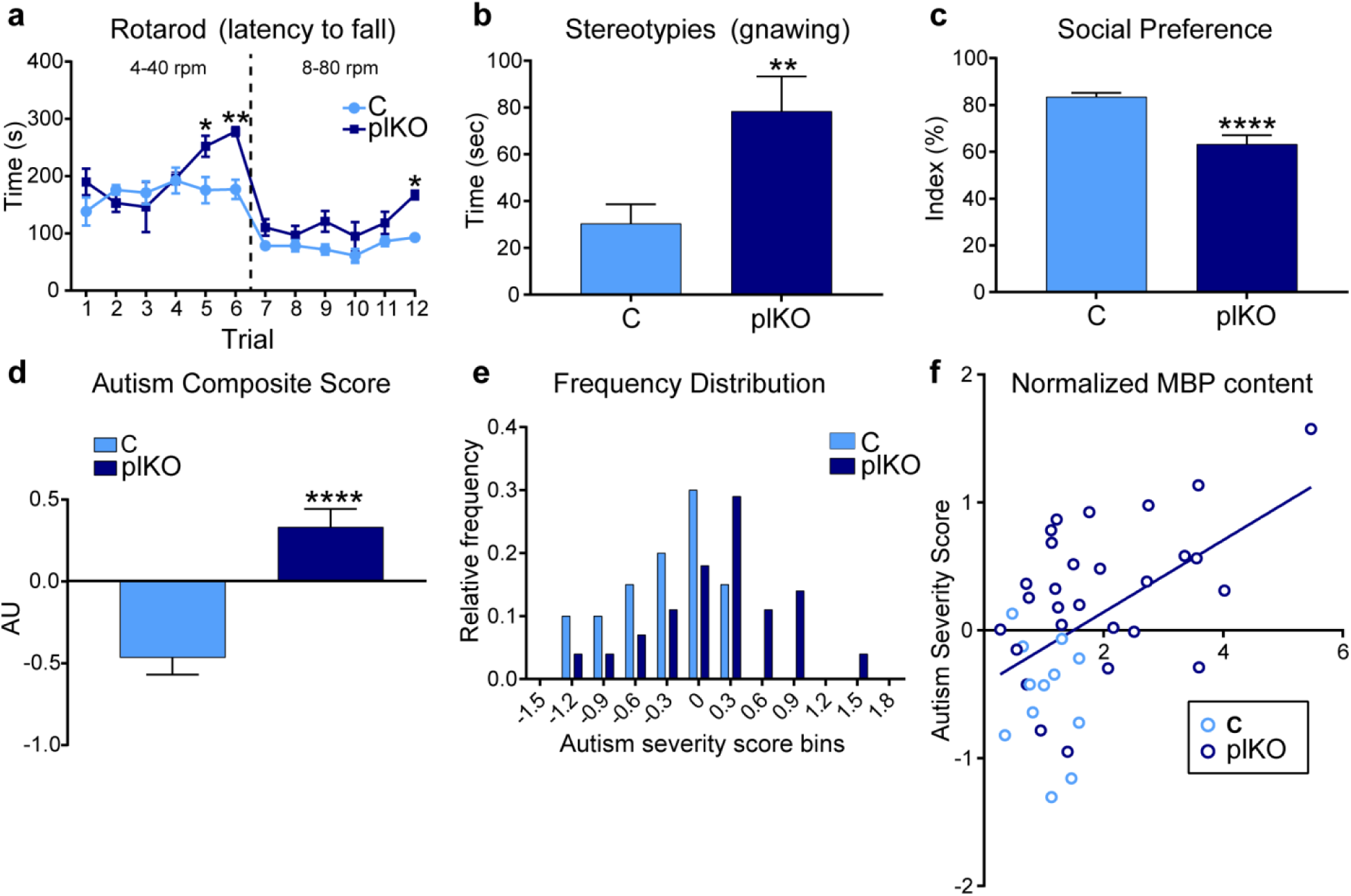
Male plKO mice exhibit ASD-like behaviour at P30. **a.** Accelerating rotarod performance in males (n=4-6/group). Data presented as means ± SEM; *p<0.05, **p<0.01 (two-way repeated-measures ANOVA with Sidak’s multiple comparisons test). **b.** Spontaneous repetitive behaviour (stereotypies) over 15 min (n=20-28/group). Data presented as means ± SEM; **p<0.01 (two-tailed unpaired Student’s t test with Welch’s correction). **c.** 3-chamber sociability test (n=20-28/group). Data presented as means ± SEM; ****p<0.0001 (two-tailed unpaired Student’s t test with Welch’s correction). **d.** ASD composite severity score (n=20-28/group). Data presented as means ± SEM; ****p<0.0001 (two-tailed unpaired Student’s t test with Welch’s correction. **e.** Relative frequency distribution of autism composite score bins (n=20-28/group). **f.** Positive correlation between cerebellum MBP levels determined by Western blot (normalized with GAPDH) and autism severity score (n=12C and 28plKO). Linear regression (R^2^=0.2548, F=12.99). Deviation from zero: p=0.0009.

Since cerebellum is a major cognitive structure that is often associated with ASD symptoms in humans and genetic mouse models^17,18^, we next asked whether mice with placental ALLO insufficiency show autistic-like features in adolescence. We first investigated stereotyped motor behaviour in spontaneously behaving mice. Male, but not female, plKO mice exhibited specific motor stereotypies, particularly increased gnawing (Fig. 3b; Extended Data Fig. 9g). On the other hand, male plKO mice spent less time digging spontaneously or during a marble burying test (Extended Data Fig. 9e,f). Increased stride length in humans and mice, as well as decreased digging in mice, have been noted as secondary ASD-like features^19-22^, particularly in males. Social behaviour, assessed using the 3-chamber sociability test, revealed significant social interaction deficits in plKO males only (Fig. 3c; Extended Data Fig. 9h), also consistent with an ASD-like behavioural phenotype. ASD is characterized by its male predominance, and stereotyped behaviour and impaired sociability represent two key hallmarks of ASD^23^, suggesting that placental hormone insufficiency may play a causal role in later developmental neuropsychiatric disease.

We determined an autism severity composite index based on previously established scoring systems^24^. A z-standardization of single behavioural readouts (including repetitive behaviour and sociability; Extended Data Fig. 9i) was performed, such that higher values represent higher symptom severity^24^. The average z-score was significantly higher in plKO males (Fig. 3d). The relative frequency distribution of individual scores revealed two distinct Gaussian distribution curves, with that of plKO mice shifted to the right (Fig. 3e). Interestingly, some of the plKO mice exhibited scores similar to those of control mice, suggesting a spectrum of behavioural changes in male plKOs. This variability may arise from natural variations in placental progesterone and ALLO levels, variable deletion of *Akr1c14*, mixed genetic background of the mice, or subtle changes in hormone exposure due to *in utero* position of the foetus.

Using MBP levels as a proxy for myelination, we found a significant positive correlation cerebellar myelination and autism symptom severity in plKO males (Fig. 3f). This finding is of particular interest knowing that there is transient WM hyperplasia and increased expression of myelin markers in the cerebellum of ASD patients, especially at young ages^6,7^, as it suggests that MBP could be used as a potential marker for ASD. Overall our mouse model recapitulates many characteristics of epidemiological studies, including the greater male vulnerability to perinatal brain injuries and ASD^7,16,21,25-28^.

The therapeutic potential of ALLO to rescue cerebellar WM and behavioural changes was then tested by administering ALLO (10 mg/kg) or vehicle (sesame oil) to dams carrying mixed litters at E15.5 to mimic the timing of the endogenous murine ALLO peak. A single injection of ALLO during gestation prevented the increase in MBP, normalized OL density in the cerebellar WM, and reduced the autism severity score in ALLO-treated plKO males compared to vehicle-treated plKO (Fig. 4). Interestingly, C mice exposed to ALLO exhibited decreased social preference (Extended Data Fig. 9j), suggesting that excess ALLO exposure during foetal life might be detrimental. Our results support the potential therapeutic utility of ALLO administration during gestation if ALLO or its precursors are determined to be low (as might occur with chronic placental insufficiency), but additionally suggest the need to maintain foetal ALLO exposure within a defined physiological window.

**Figure 4.**
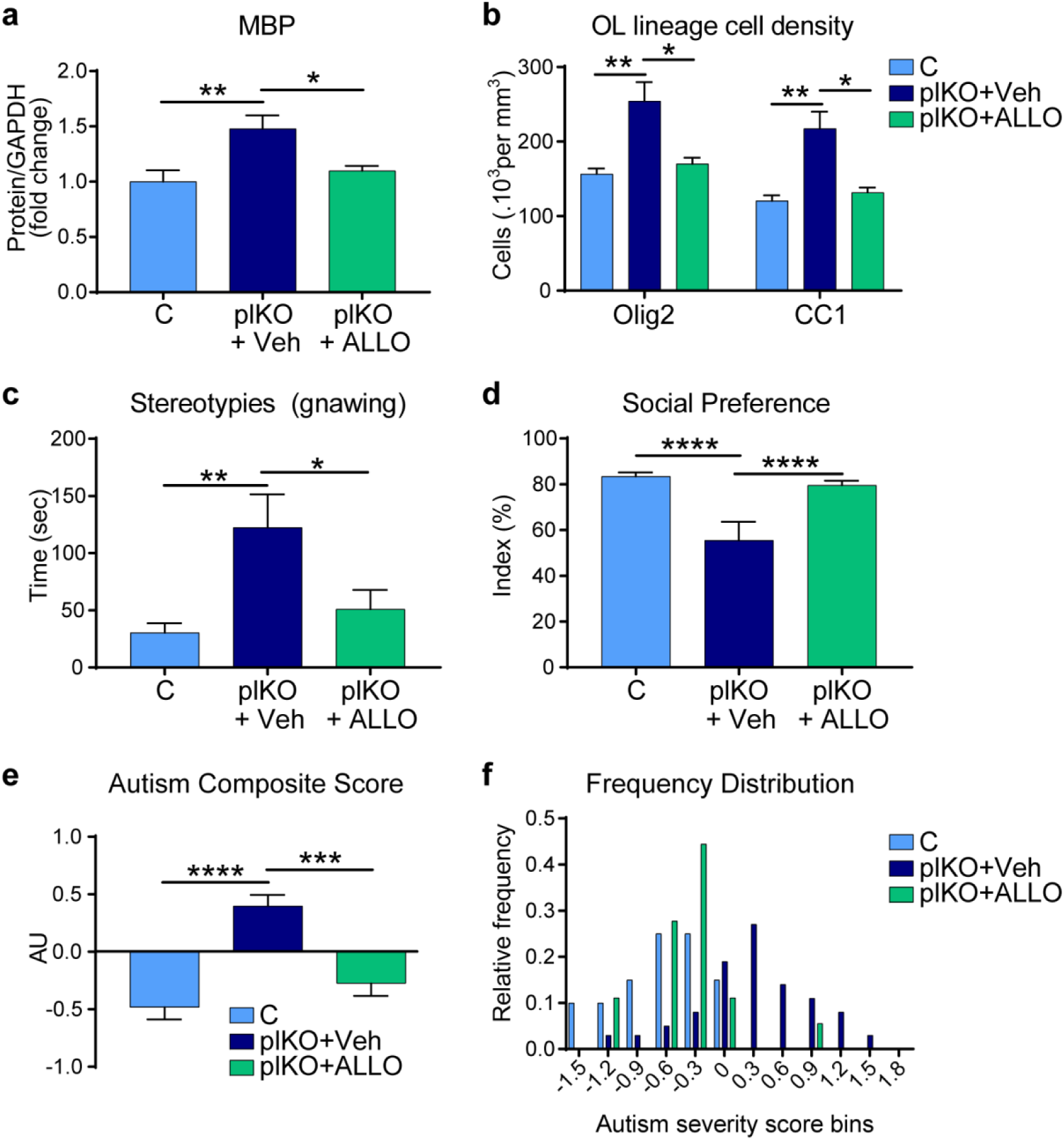
ALLO administration at E15.5 rescues cerebellar WM and behavioural impairments in male plKO mice at P30. **a.** Western blot determination of MBP contents in the cerebellum. Normalized data to GAPDH contents is presented as mean fold changes ± SEM (n=6-8/group). *p<0.05; **p<0.01 (one-way ANOVA with Tukey’s multiple comparisons test). **b.** Quantification of Olig2+ and CC1+ cell densities within the cerebellar WM (n=7C, 8plKO+Veh and 4plKO+ALLO). Data presented as means ± SEM. *p<0.05; **p<0.01 (two-way ANOVA with Sidak’s multiple comparisons test). **c.** Stereotypies (gnawing) time over 15 min (n=20C, 8plKO+Veh, and 17plKO+ALLO). Data represented as means SEM; *p<0.05; **p<0.01 (one-way ANOVA with Tukey’s multiple comparisons test). **d.** 3-chamber sociability test (n=20C, 8plKO+Veh, and 17plKO+ALLO). Data presented as means ± SEM; ****p<0.0005 (one-way ANOVA with Tukey’s multiple comparisons test). **e.** ASD composite score n=20C, 36plKO(+Veh) and 17plKO+ALLO. ***p<0.005; ****p<0.0005 significant difference between groups (one-way ANOVA with Tukey’s multiple comparisons test). plKO and plKO+Veh mice were combined since there was no significant difference between both groups. **f.** Relative frequency distribution of autism composite score bins. plKO and plKO+Veh mice were combined (n=20C, 36plKO (+Veh), 17plKO+ALLO).

Our study empirically defines a critical role for a placental hormone that can alter foetal brain development in late gestation with long-term functional consequences. Many neuroactive hormones are produced in the placenta in late gestation that may impact the late stages of neuro- and gliogenesis, neuronal migration and circuit formation. We focused on ALLO because this placental steroid hormone normally peaks in the second half of gestation, when placental insufficiency becomes evident and when preterm birth occurs. Our experiments demonstrate a treatment-reversible link between specific placental ALLO insufficiency and sex-specific cerebellar hypermyelination associated with an ASD-like phenotype. Increased risk of ASD diagnosis following extremely preterm birth^29^, particularly in males, suggest that loss of placental hormones may contribute to specific human neurobehavioural outcomes in previously unanticipated ways. Our findings lay the groundwork for developing hormone replacement strategies to normalize the developmental milieu and prevent long-term impairments in neurobehaviour.

## METHODS

### Mice

All the procedures on experimental animals were performed in accordance with the protocol approved by the Institutional Animal Care and Use Committee at Children’s National Medical Center (protocol # 30534). Mice were housed in a 12:12 light:dark cycle, and had *ad libitum* access to food and water.

The Akr1c14-floxed mouse was constructed with inGenious Targeting Laboratories (Ronkonkoma, NY). The construct targeted a 3.15kb region that includes exons 7-9 of *Akr1c14*, with a 5’ homology arm extending to the position of a LoxP-FRT flanked Neo cassette and a 3’ homology arm extending to a single LoxP site just 5’ of exon 10 (Fig. 1c). The targeting vector was electroporated into embryonic stem cells, which were cloned and screened. Confirmed clones were micro-injected into C57Bl/6 blastocysts. The Neo cassette was later removed using flipase. The resulting mice are designated as the *Akr1c14*-floxed line. Akr1c14^fl/wt^ and Akr1c14^fl/fl^ mice appear phenotypically normal. The final placental 3αHSD knockout was generated by successive crosses of progenitor Akr1c14-floxed mice and Cyp19a-Cre mice (generous gift of G. Leone, Iowa), which express Cre exclusively in placental trophoblast cells^9^ (Fig. 1c; Extended Data Fig. 2a-d). Mouse, unlike humans, does not express endogenous placental Cyp19a, which codes for aromatase, but the human Cyp19a promotor effectively drives placenta-specific Cre expression^9^. Cyp19a-Cre females were crossed to homozygous Akr1c14^fl/fl^ males. Female offspring of these crosses that were positive for Cre with two floxed alleles were used as the dams of experimental mice. These Akr1c14^fl/fl^ females carrying Cyp19a-Cre were crossed to Akr1c14^fl/fl^ males, so that 1/2 of resulting offspring are both homozygous Akr1c14^fl/fl^ and positive for placental Cre. These mice in which *Akr1c14* is deleted specifically in the placenta are Akr1c14^Cyp19a-/-^, designated as Akr1c14^Cyp19a^KO (plKO) for simplicity. An additional 1/2 of offspring are negative for placental Cre and designated as controls (C).

### Drug injections

Pregnant dams received one intraperitoneal (i.p.) injection of allopregnanolone (Tocris Bioscience, Minneapolis, MN, USA; #3653) diluted in sesame oil (vehicle; Sigma-Aldrich, Saint-Louis, MO, USA; #S3547) at E15.5 at a dose of 10 mg/kg of body weight. BrdU (50 mg/kg) dissolved in saline solution was injected i.p. in dams at E15.5.

### Mass Spectrometry

#### Steroid extraction

Pregnenolone (PREG), progesterone (PROG), allopregnanolone (ALLO), epiallopregnanolone (EPIALLO), allotetrahydrodeoxycorticosterone (ALLOTHDOC) and 3α5α-tetrahydrotestosterone (3α5α-THT) were identified and quantified simultaneously in individual tissues by GC/MS/MS as previously described ^30^. Placental (65 – 185 mg) and foetal brain (61 – 130 mg) tissues of male and female WT and plKO mice at E17.5 were weighed and stored at −20 °C until GC/MS/MS analysis. Briefly, steroids were first extracted from placentas and brains with 10 vols of methanol (MeOH) and the following internal standards were added to the extracts for steroid quantification: 2 ng of epietiocholanolone for PREG, ALLO, EPIALLO, ALLOTHDOC and 3α5α-THT, 2 ng of ^13^C_3_-PROG for PROG. Samples were purified and fractionated by solid-phase extraction with the recycling procedure ^31^. The extracts were dissolved in 1 ml MeOH and applied to the C18 cartridge (500 mg, 6 ml, International Sorbent Technology, IST), followed by 5 ml of MeOH/H2O (85/15, vol/vol). The flow-through, containing the free steroids, was collected and dried. After a previous re-conditioning of the same cartridge with 5 ml H2O, the dried samples were dissolved in MeOH/H2O (2/8, vol/vol) and re-applied. The cartridge was then washed with 5 ml H2O and 5 ml MeOH/H2O (1/1, vol/vol) and unconjugated steroids were eluted with 5 ml MeOH/H2O (9/1, vol/vol). The fraction containing the unconjugated steroids was then filtered and further purified by high performance liquid chromatography (HPLC). The HPLC system is composed of a WPS-3000SL analytical autosampler and a LPG-3400SD quaternary pump gradient coupled with a SR-3000 fraction collector (Thermo Fisher Scientific, Waltham, MA, USA). The HPLC separation was achieved with a Lichrosorb Diol column (25 cm, 4.6 mm, 5 μm) in a thermostated block at 30 °C. The column was equilibrated in a solvent system of 90% heptane and 10% of a mixture composed of heptane/isopropanol (85/15, vol/vol). Elution was performed at a flow-rate of 1 ml/min, first 90% heptane and 10% of heptane/isopropanol (85/15, vol/vol) for 15 min, then with a linear gradient to 100% acetone in 2 min. The column was washed with acetone for 15 min. Steroids were collected in the time range 15 – 29 min, and were derivatized with 25 μl of HFBA and 25 μl of anhydrous acetone for 1 h at 20 °C. Samples were dried under a stream of nitrogen and resuspended in heptane.

#### GC/MS/MS analysis

GC/MS/MS analysis of the biological extracts was performed using an AI 1310 autosampler, a Trace 1310 gas chromatograph (GC), and a TSQ 8000 mass spectrometer (MS) (Thermo Fisher Scientific). Injection was performed in the splitless mode at 250 °C (1 min of splitless time) and the temperature of the gas chromatograph oven was initially maintained at 80 °C for 1 min and ramped between 80–200 °C at 20 °C/min, then ramped to 300 °C at 5°C/min and finally ramped to 350 °C at 30 °C/min. The helium carrier gas flow was maintained constant at 1 ml/min during the analysis. The transfer line and ionization chamber temperatures were 300 °C and 200 °C, respectively. Electron impact ionization was used for mass spectrometry with ionization energy of 70 eV and GC/MS/MS analysis was performed in multiple reaction monitoring (MRM) mode with Argon as the collision gas. GC/MS/MS signals were evaluated using a computer workstation by means of the software Excalibur®, release 3.0 (Thermo Fisher Scientific). Identification of steroids was supported by their retention time and two or three transitions. Quantification was performed according to the more abundant transition with a previously established calibration curve. The range of the limit of detection was roughly 0.5–20 pg according to the steroid structure. The GC/MS/MS analytical procedure was fully validated in terms of accuracy, reproducibility and linearity in mouse brain ^30^.

### Behavioural testing

Tests were performed on 30- to 37-day old mice. No more than 3 behavioural tests were done on each mouse, accordingly to our IACUC recommendations. A resting period was allowed between 2 consecutive tests.

#### Rotarod

The accelerating rotarod performance test is a standard rodent assay for motor-associated functions such as coordination and balance. A 4 day testing paradigm (3 sessions per day) was performed as previously described^16^ with a continuously accelerating rod that forces animals to modulate their balance. For the first 2 days, animals were placed on the rod which accelerated from 4-40 rpm in 180 seconds. The test was stopped when the animal fell off the rod, was unable to maintain coordinated movements, or 180 seconds passed. Additionally, if animals displayed “cartwheeling”, or clutching the rod and circling without walking, the test was stopped. To increase the challenge, for days 3-4 the acceleration of the rod increased from 8-80 rpm in 180 seconds. The latency to fall from the rod, and terminal velocity was then recorded. Linear regression analysis on data from each individual mouse was used to estimate initial motor coordination (given by the intercept) and learning rate (given by the slope).

#### Spontaneous behaviour/stereotypies

Further assessment of repetitive behaviour was done to test autism-like behaviour. Mice were allowed to habituate to testing room before placement in a novel, empty cage with bedding materials. A video camera was set with a lateral view to record spontaneous behaviour for 15 minutes. Videos were manually scored for time spent engaging in behaviours including digging, gnawing, grooming, rearing and jumping.

#### Socialization test

The three-chamber test (Crawley’s paradigm^32^) was used to assess social behaviour as previously described^23^. A three-chamber Plexiglas box was used with small openings in each dividing wall to allow free access to each chamber. The centre chamber was kept empty, while the flanking chambers were each equipped with an identical wire cup. After a 10 minute habituation period to allow for exploration of the 3-chamber equipment (session 1), an unfamiliar adult mouse of similar weight and coloration was placed within the wire cup in one of the side chambers and an “object” within the wire cup in the opposite chamber. The dividers were raised to allow the test subject to move freely throughout all three chambers of the apparatus over a 10-minute test session (session 2). The second session was analysed for social preference given by the social preference index (SPI). SPI was calculated as follows: if time spent with the mouse (social) is S and the object (non-social) is NS, then SPI=S/(S+NS).

#### Determination of the autism composite severity score

We used the same procedure as previously described by ^24^. The score calculation allows for the combination of individual discrete symptoms of the autistic syndrome, in a continuous manner, so that higher values reflect higher severity of autistic-like behaviour. For each mouse, we recorded their spontaneous behaviours (grooming, digging, rearing, jumping and gnawing) and determined their social preference index in the 3-chamber test. The significant readouts (digging time, gnawing time and social preference index) were then z-standardized and averaged to establish the autism composite severity score.

### Histology

#### Section preparation and staining

Animals were anesthetized using isoflurane (Isothesia, Henry Schein Animal Health, Dublin, OH, USA) and transcardiacally perfused with 20 mL 1×PBS followed by 30 mL 4% paraformaldehyde (PFA), as previously described. Brains were post-fixed for 24h in 4% paraformaldehyde and cryoprotected in 20% sucrose in PBS. Serial 40-μm-thick sagittal brain/cerebellar sections were then collected using a sliding microtome. Placentas were collected at E17.5 after deep anaesthesia of the dams, drop-fixed in 4% PFA and cryoprotected in 30% sucrose. Frozen 20-μm cross-sections were obtained with a cryostat. Immunohistochemistry on brain or placenta sections was performed using the following antibodies: rabbit anti-MBP (Abcam, Cambridge, MA, USA; #ab40390; 1:500), mouse anti-APC clone CC1 (EMD Millipore, Burlington, NA, USA; #MABC20; 1:500), rabbit anti-Olig2 (Abcam; #ab9610, 1:500), rabbit anti-MBP (Abcam; #ab40390; 1:500), mouse anti-neuronal nuclei (NeuN) (Millipore; #MAB377; 1:500), goat anti-NeuroD1 (RD Systems, Minneapolis, MN, USA; #AF2746; 1:500), rabbit anti-calbindin (Swant, Marly, Switzerland; #CB38, 1:1000), rat anti-mouse PDGFRα (CD140a; BD Biosciences, San Jose, CA, USA; 1:500), chicken anti-GFP (Abcam; #ab13970; 1:500), rabbit anti-MCT4 (SLC1A3; ABclonal, Woburn, MA, USA; #A15722; 1:200). Sections were incubated with primary antibodies overnight at 4°C containing 0.3% Triton and 10% normal donkey serum. Sections were then incubated with secondary antibodies (1:500) together with DAPI (Invitrogen, Carlsbad, CA, USA; #D1306: 1:1000) for 2 hours at room temperature. FluoroMyelin™ Green (Invitrogen; #F34651; 1:500) was also added at this time for those sections. Secondary antibodies were used as follows: donkey anti-mouse Alexa-488 (Invitrogen; #A-21202), donkey anti-rabbit Alexa-488 (Invitrogen; #A-21206), donkey anti-mouse Alexa-555 (Invitrogen; #A-31570), donkey anti-rabbit Alexa-555 (Invitrogen; #A-31572), donkey anti-mouse Alexa-647 (Invitrogen; #A-31571), donkey anti-rabbit Alexa-647 (Invitrogen; #A-31573). Finally, floating sections were mounted and cover slipped using ProLong™ Gold Antifade Mountant (Thermo Fisher Scientific; #P36930) before imaging.

#### Fluorescence imaging and Cell Counting

Whole brain or cerebellum sagittal sections were imaged using a Virtual Slide Microscope (VS 120, Olympus Life Science, Waltham; MA, USA) under a 20× objective. High magnification images were taken under a confocal microscope (TCS SP8, Leica Microsystems, Buffalo Grove, IL, USA). Z-stack images were acquired with a step size of 1.55 μm and viewed using NIH ImageJ (http://imagej.nih.gov/ij). Cell quantification was done on 6 sections per animal, homogeneously distributed throughout the cerebellar vermis, with 6-7 animals per group. The different cerebellar layers were delineated manually using the freehand selection tool in ImageJ. The density of Olig2-, CC1- and BrdU-positive cells was estimated using Stereo Investigator® (MBF Bioscience, Williston, VT, USA). After delineating the cerebellar WM using the freehand selection tool on corresponding DAPI images, immunopositive cells were manually counted under the 40x objective (Zeiss Axio Imager M2, Thornwood, NY, USA) using the optical fractional as a probe in grids randomly distributed on and covering 30% of the cerebellar WM. Cell densities were then calculated by using the individual WM area values and the mean measured section thickness. The linear density of Purkinje cells (Purkinje cells per mm) was determined in the lobule VI-VII by drawing a freehand line through the centre of the cell bodies.

### Electron microscopy

#### Sample preparation

P30 mice were perfused under deep isoflurane anaesthesia with 20 mL 0.12M cacodylate buffer followed by 30 mL fixative made of 2.5% glutaraldehyde and 1% paraformaldehyde (electron microscopy grade, EMS #16221 and #15713) in cacodylate buffer (pH7.4). Brains were removed and post-fixed for 1 hour in the same fixative. 300-μm-thick sagittal slices were then made using a vibratome (VT 1000S, Leica Biosystem, Buffalo Grove, IL, USA). Slices were post-fixed with 1% osmium tetroxide in 0.12 M cacodylate buffer for 2 hours at room temperature and “en block” stained with 1% uranyl acetate in 0.1M acetate buffer overnight at room temperature. After several washes in acetate buffer, slices were dehydrated by passing them through increasing concentrations of ethyl alcohol (up to 100%). Slices were then progressively infiltrated in epoxy resin, and placed 48h at 60°C for resin polymerization. 100-nm-thick sagittal ultrasections were performed using an ultramicrotome (Ultracut UC7, Leica Biosystem) through the whole cerebellum (+brainstem) or the whole anterior brain (for corpus callosum analyses).

#### Image acquisition

Ultrathin sections were observed with an FEI Helios NanoLab 660 FIBSEM field emission scanning electron microscope (FEI, ThermoFischer Scientific) using extreme high resolution and equipped with a concentric (insertable) higher-energy electron detector. Low-magnification images (600×) were first taken in order to delineate our regions of interest (vermal inter-lobule 6-7 of the cerebellum or mid-body corpus callosum). Then high magnifications images of our regions of interest (20,000-50,000×) were taken using 4 kV and 0.2 current landing voltage.

#### Myelin thickness measurement

The outer (including myelin sheath) and inner diameter of 100 randomly selected myelinated axons was measured on one ultrathin section per animal (3 animals/group). The g-ratio (equal to the ratio of the inner-to-outer diameter of a myelinated axon) of each axon was calculated. Scatter plots of the axon calibre and g-ratios were then analysed. The slopes of the linear regressions from C and plKO mice were compared.

### Western blots

Whole cerebellums were dissected out and homogenized in 300 μL radioimmunoprecipitation assay (RIPA) lysis buffer consisting of (in mM) 50 Tris-HCl, pH 7.4, 150 NaCl, 2 EDTA, 50 NaF, 1 Na3VO4, 1% Triton X-100, 0.1% SDS, 0.5% Na-deoxycholate, and a Protease/PhosphataseInhibitor Cocktail (Santa Cruz Biotechnology, Santa Cruz, CA, USA). Following centrifugation at 14,000g for 10 min, protein concentration was determined using Bradford protein assay kit (Bio-Rad, Hercules, CA, USA). Sample total proteins were resolved by sodium dodecyl sulfate-polyacrylamide gel electrophoresis using 10% Bis-Tris precast gel (Thermo Fisher Scientific) and transferred to polyvinylidene fluoride (PVDF) membranes. Membranes were incubated with blocking buffer consisting of 4% non-fat milk in 1% Tween-20 in Tris buffered saline (TBS-T) for 1h, followed by overnight incubation at 4°C with one of the following primary antibodies diluted in 3% BSA TBST-T: rabbit anti-MBP (Abcam; #ab40390; 1:500), rabbit anti-MAG (Thermo Fisher Scientific; #PA5-79620; 1:500), rabbit anti-MOG (Abcam; 1:500), mouse anti-cofilin-1 (Cfl1) (Santa Cruz Biotechnology, Dallas, TX, USA; #sc-53934; 1:500), mouse anti-prosaposin (PSAP) (Santa Cruz Biotechnology; #sc-390184; 1:500), rabbit anti-hnRNPκ (R332) (Cell Signaling Technology; #4675; 1:500) and rabbit anti-GAPDH (Cell Signaling Technology, Danvers, MA, USA; #5174; 1:2000). After three washes with TBS-T, membranes were incubated with horseradish peroxide (HRP)-conjugated secondary antibodies (Jackson ImmunoResearch, West Grove, PA, USA; 1:2000), and protein bands were visualized using chemiluminescent ECL detection system (Bio-Rad) according to manufacturer’s instruction. Signal intensities of protein bands were quantified using Image J software (http://rsb.info.nih.gov/ij/) and normalized with GAPDH as an internal control. When more than 10 samples, and thus several membranes, needed to be compared at once, 2 common samples resolved in the different electrophoresis gels, and used as reference for quantification.

### Statistics

Mouse assignment to groups was not randomized, since it relies on their known genotype (C vs plKO), but mice from at least three different litters were used in each group for all experiments. No litter-associated effect was seen. All data analyses were conducted blind to group allocation. Statistical analysis was carried out with GraphPad Prism 7 software. The sample sizes are shown in the legends and chosen to meet or exceed sample sizes typically used in the field. ROUT methodology with Q value equal to 1 was utilized to determine outliers. Prior to conducting all analyses, variable distributions were evaluated for normality and transformed if needed. If transformation was not possible, non-parametric methods were used. Differential gene expression across development in C and plKO mice was analysed using one-way ANOVA with Dunnett’s multiple comparisons test. Comparisons between two groups were analysed using parametric test (unpaired t-test with Welch’s correction) or non-parametric test (Mann-Whitney test). Analyses involving data from three or more groups whilst considering only one independent variable were performed using one-way ANOVA with Tukey’s multiple comparisons test. Two-way ANOVA was used to compare means across two or more dependent variables. Two-way repeated measures ANOVA with Sidak’s multiple comparisons tests were used for the Erasmus ladder and the Rotarod. Summary data are presented in the text as mean ± SEM from n animals. Differences were considered significant at p < 0.05. A supplementary statistics methods checklist is available.

## Acknowledgements

We thank Dr. G. Leone for providing us Cyp19a-Cre mice, Drs. Li-Jin Chew and Ioannis Papazoglou for comments on the manuscript, Dr. Joseph Scafidi for discussion and Sarah E. P. Thau for scoring the beam walking test. This work was funded by the National Institutes of Health R01HD092593 and 3R01HD092593-S1 (A.P.) and R37NS109478 (V.G.), the Simons Foundation (SFARI Pilot Award; A.P.), the Children’s National Board of Visitors (A.P. and V.G.), and the Research Foundation of Cerebral Palsy Alliance (#3720; A.P.).

## Author contributions

C.-M.V. and A.A.P. conceived the project and designed the experiments. C.-M.V. and J.S. performed pilot experiments. J.O.R., H.L. and D.B. performed the genetic validation of the model. Y.I., K.H.-T., T.S. and C.-M.V. performed the RNA sequencing analysis. H.L. performed the qRT-PCRs. C.-M.V. and J.S. performed the histological experiments; C.-M.V. did the analyses. C.-M.V., J.S. and S.S. performed the behavioural experiments. A.S. and V.G. contributed to the Erasmus ladder data acquisition and analysis. C.-M.V., C.C.-P. and A.P. performed the electron microscopy experiments. C.-M.V. performed the biochemical analyses. P.L. and M.S. performed the mass spectrometry experiments. S.S. and J.S. performed the genotyping. C.-M.V. and A.A.P. wrote the manuscript. All authors reviewed and revised the manuscript.

## Data availability

Raw and processed data are available from the corresponding authors upon request.

## SUPPLEMENTAL MATERIAL

### Methods

#### *In situ* hybridization

For analysis of *Akr1c14* mRNA, E17.5 placental tissue was drop fixed overnight in 4% PFA and then transferred through a sucrose gradient of 10%, 20% and 30% over three days. Tissue was sectioned at 12 µm and assayed for *Akr1c14* RNA using the Advanced Cell Diagnostics (Newark, CA) RNAScope system and an ACD probe designed to target the 621-1989 region of *Akr1c14* (NM_134072.1).

#### 3’mRNA-sequencing

##### Tissue collection and RNA extraction

Cerebellum, bilateral cerebral cortices and bilateral hippocampi from C and plKO mice (males and females) at postnatal day (P) 30 (3 animals/group) were dissected and flash frozen. Tissue homogenization and total RNA extraction were performed using the *mir*Vana Isolation kit (Thermo Fisher Scientific; #AM1560) according to the manufacturer’s instructions.

##### Library Preparation and Sequencing for mRNA

The cDNA libraries were prepared using the QuantSeq 3’mRNA-Seq Library Prep Kit FWD for Illumina (Lexogen, Greenland, NH, USA) as per the manufacturer’s instructions. Briefly, total RNA was reverse transcribed using oligo (dT) primers. The second cDNA strand was synthesized by random priming, in a manner that DNA polymerase is efficiently stopped when reaching the next hybridized random primer, therefore only the fragment closed to the 3’ end was captured for indexed adapter ligation and PCR amplification. The processed libraries were assessed for its size distribution and concentration using BioAnalyzer High Sensitivity DNA Kit (Agilent Technologies, Santa Clara, CA, USA; #5067-4626). Pooled libraries were diluted to 2 nM in EB buffer (Qiagen, Redwood City, CA, USA; #19086) and then denatured using the Illumina protocol. The libraries were pooled and diluted to 3 nM using 10 mM Tris-HCl, pH 8.5 and then denatured using the Illumina protocol. The denatured libraries were diluted to 10 pM by pre-chilled hybridization buffer and loaded onto a TruSeq v2 Rapid flow cell on an Illumina HiSeq 2500 and run for 50 cycles using a single-read recipe according to the manufacturer’s instructions. De-multiplexed sequencing reads were generated using Illumina bcl2fastq (released version 2.18.0.12) allowing no mismatches in the index read.

##### Data Analysis

After the quality and polyA trimming by BBDuk^1^ and alignment by HISAT2 (version 2.1.0)^2^, read counts were calculated using HTSeq^3^ by supplementing Ensembl gene annotation (GRCm38.78). DE analysis and M-A plots were done using TCC R package (version 1.12.1, https://bioconductor.riken.jp/packages/3.3/bioc/manuals/TCC/man/TCC.pdf)^4^. After getting differential expressed genes (DEGs), hierarchical clustering of DEGs as a heatmap was performed using Partek Genomics Suite (version 6.6, http://www.partek.com/partek-genomics-suite/) (Partek). No outlier was identified in our biological replicates, as indicated by principal component analysis (PCA) and scatter matrices. Differentially expressed genes (DEGs) met the following guidelines: p < 0.05; fragments per kilobase of transcript per million mapped reads (FPKM) values >1; and fold change (FC) >1.5. The limits of these cutoffs were validated by RT-PCR on samples from other mouse cohorts. Ingenuity Pathway Analysis (IPA; Qiagen, Redwood City, CA) was used to identify the top biological functions and disease processes that were differentially regulated. This software is based on the ingenuity pathway knowledge base (IPKB) for genetic interaction, which derives from the scientific literature, each network connection being supported by previous publications ^5^. To determine and visualize the degree of gene overlaps in datasets, Venn analysis was performed using Venny 2.1 (http://bioinfogp.cnb.csic.es/tools/venny/).

#### Real-Time PCR (RT-PCR)

Tissues were homogenized in TRIzol™ Reagent (Thermo Fisher Scientific); total RNA was extracted with the RNeasy Mini Kit (Qiagen, Venlo, Netherlands; #74104).1 µg of RNA was used to make cDNA with the iScript cDNA Synthesis Kit (Bio-Rad, Hercules, CA, USA; #1708891). All primer pairs were designed and validated in-house for efficiency and specificity. RT-PCR experiments were performed on cDNA samples in presence of PowerUp SYBR Green Master Mix (Thermo Fisher Scientific; #1725271) with specific primers at 100 nM using the CFX96 Touch™ Real-Time PCR Detection System (Bio-Rad). The cDNA-generated signals for target genes were internally corrected with phosphoglycerate kinase 1 (PGK1). The regulation was determined with the 2-ΔΔCq method^6^.

#### Erasmus ladder

The Erasmus ladder (Noldus, Leesburg, VA, USA) is a fully automated system allowing for the assessment of baseline gait and locomotor coordination, and associative cerebellar learning. The instrument consists of a horizontal ladder with 2×37 pressure sensitive rungs for both the left and right sides, stationed between two shelter boxes. Each shelter is equipped with a white LED spotlight and pressurized air outlet serving as cues to signal time to departure from shelters. As the mouse is prompted to move, pressure sensors continuously monitor and analyse the walking pattern of the mouse in real time, sending data to the computer system, which in turn adjusts air pressure, calculates interventions and predicts future steps. Through this prediction, the software can also be programmed to move rungs mid-trial by high speed pneumatic slides to create an obstacle. Mice were trained with 42 runs per day for 4 successive days without obstacles. Missteps, trial times and jumps were measured for each run. Following training, cerebellar associative motor learning was assessed on days 4-8 with the introduction of a tone as a conditioning stimulus (CS) and a rising rung (obstacle) as the unconditioned stimulus (US). These perturbations were used to evaluate mouse ability to adapt motor behaviour in real time. Repeated perturbations were used to assess associative motor learning throughout successive trials. With the inter-stimulus interval fixed at 285 ms, mice that associated the tone with the obstacle learned to increase walking speed to avoid being hit. In this sense they decreased their “pre-perturbation step time” (step prior to US) and their “post-perturbation step time” (step just after the US). The Erasmus ladder is particularly valuable for detailed phenotyping of deficits associated with cerebellar pathology ^7^.

#### Inclined beam test

The inclined beam walking task was performed to assess balance and gait on ∼P30 mice as previously described^8,9^. A wooden beam, 1 meter in length and 1cm in width was set at a 30-degree angle leading to a darkened box as a finishing point. To encourage movement up the beam, a lamp was used to shine light above the starting point as an aversive stimulus and bedding from the home cage was placed in the goal box. The first day, mice were allowed to habituate to the beam and goal box without assessment. The next day, mice were tested 3 times on the beam, with at least 2 hours rest between each trial. Time to cross an 80 cm mark was measured and recorded by the experimenter in real time. At the base of the beam a video camera recorded the mouse from behind to assess number of foot slips with either front or hind paws during each trial.

#### Marble burying

The anxiety and stereotypic and/or obsessive-compulsive-like behaviour components of ASD-like behaviour can be assessed in mice by their behavioural responses to novel “diggable” media^10,11^ in the marble burying test. This test was performed in a clean large (26×16 cm) cage filled with 5 cm of bedding and twelve glass marbles evenly spaced on the bedding surface. The animal was left to explore the cage for 30 minutes undisturbed. Marbles covered 2/3^rds^ or more with bedding were counted as buried.

## Extended Data Figure legends

**Extended Data Figure 1.**
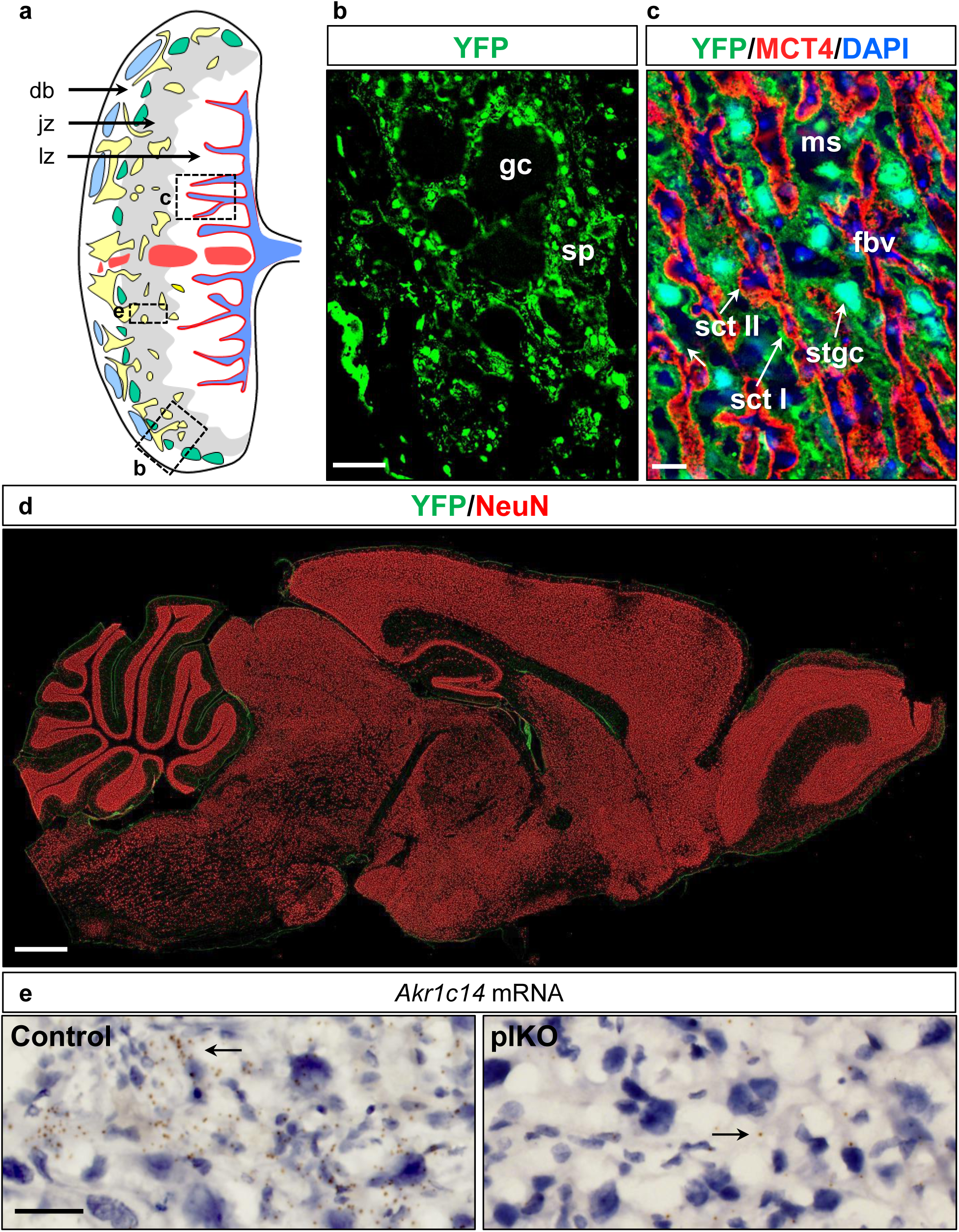
Visualization of Cyp19a driven Cre expression in the placenta of Cyp19aCre-ROSA26YFP mice. **a.** Schematic representation of a transversal section of the mouse placenta adapted from Georgiades et al., 2001 ^12^. The plane of sectioning is through the center of the placenta and perpendicular to its flat surface, such that the orientation shows maternal side on the left and the foetal side on the right. The major placental zones (db, decidua basalis; jz, junctional zone; lz, labyrinth zone) are shown; their constituent cell types are depicted by different colors. **b.** Illustration of the YFP-positive staining in the junctional zone of a mouse placenta at E17.5 in green. *Cyp19a* promoter activity (YFP+) is detected by immunofluorescence with an anti-GFP antibody in the spongiotrophoblasts (sp) surrounding glycogen cell clusters (gc). Scale bar, 100 μm. **c.** Illustration of the YFP-positive staining in the labyrinth zone of a mouse placenta at E17.5 in green. MCT4 immunostaining is used to label syncytiotrophoblasts II in red. *Cyp19a* promoter activity (YFP+) is detected in syncytiotrophoblasts I (sct I) and sinusoidal trophoblast giant cells (stgc). Scale bar, 20 μm. **d.** Sagittal section of a Cyp19aCre-ROSA26YFP mouse brain immunostained against YFP (green) and NeuN (red). No Cyp19a promotor activity was evidenced in the brain. Scale bar, 500 µm. **e**. *In situ* hybridization for *Akr1c14* mRNA in the placenta at E17.5. *Akr1c14* mRNA levels are drastically decreased in the spongiotrophoblasts (arrows) of plKO as compared with C mice. Scale bar, 15 μm. fbv, foetal blood vessels; gc, glycogen cell clusters; MCT4, monocarboxylate transporter 4; ms, maternal sinus; sct I/II, syncytiotrophoblast I/II; sp, spongiotrophoblasts; stgc, sinusoidal trophoplast giant cell.

**Extended Data Figure 2.**
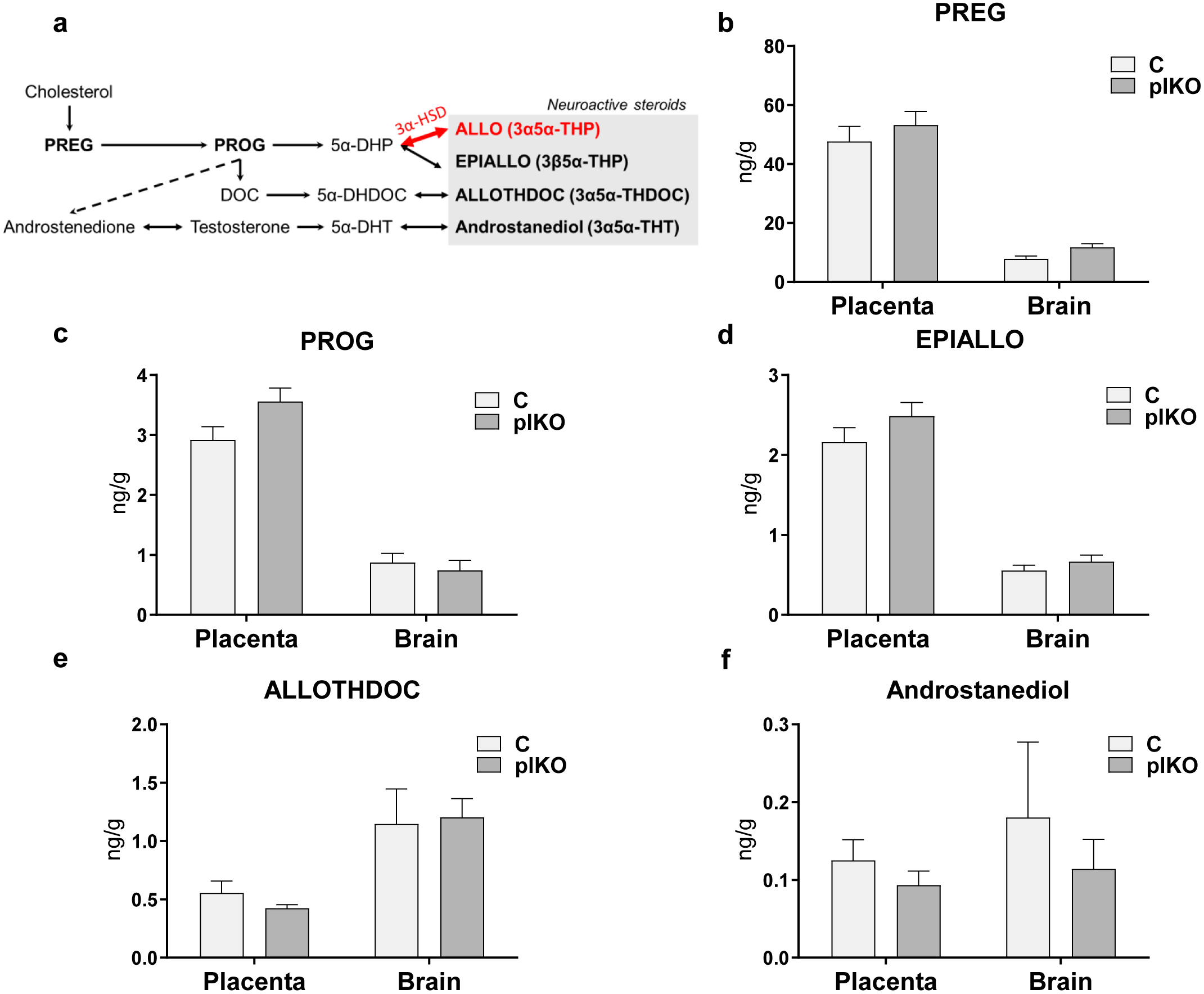
Neurosteroid synthesis pathways and chemical validation of the plKO model. **a.** Pathways from cholesterol to ALLO and other steroids known to modulate GABA-A receptors. **b-f.** The production of the neuroactive steroids other than ALLO was not impacted by the placental *Akr1c14* deletion, as confirmed by mass spectrometry. Data is presented as means ± SEM, n=9-12 pools of 3 samples/group. Two-way ANOVA with Sidak’s multiple comparisons test. 3α-HSD, 3α-hydroxysteroid dehydrogenase; 3α5α-THDOC, 3α5α-tetrahydrodeoxycorticosterone; 3α5α-THP, 3α5α-tetrahydroprogesterone; 3β5α-THP, 3β5α-tetrahydroprogesterone; 3α5α-THT, 3α5α-tetrahydrotestosterone; 5α-DHDOC, 5α-dihydrodeoxycorticosterone; 5α-DHP, 5α-dihydroprogesterone; 5α-THT, 5α-dihydrotestosterone; ALLO, allopregnanolone; DOC, deoxycorticosterone; PREG, pregnenolone; PROG, progesterone.

**Extended Data Figure 3.**
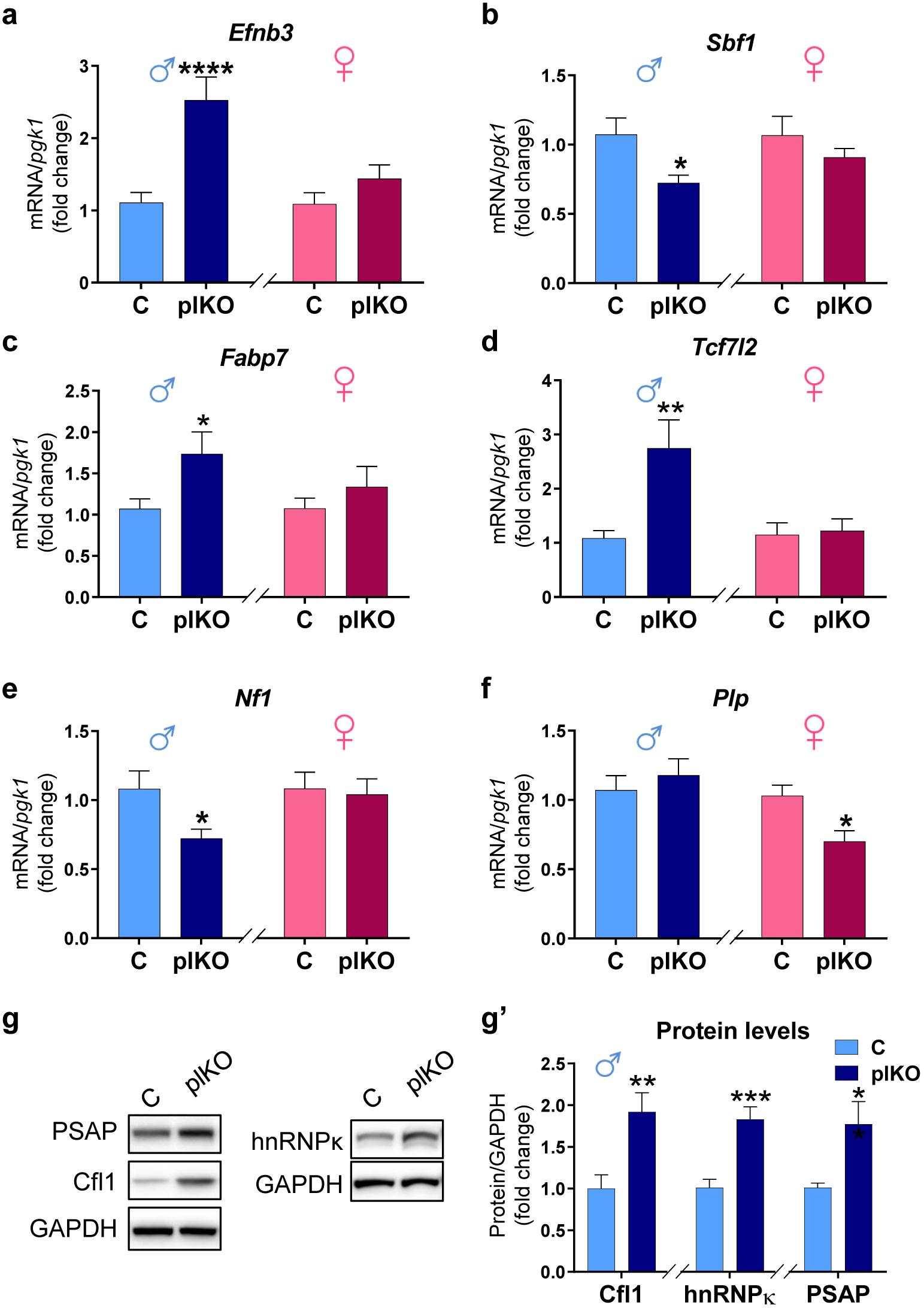
Molecular validation of the RNA sequencing cutoffs. Changes in mRNA (a-f) or protein (g-g’) contents of selected cerebellar dysregulated genes after normalization. **a.** *Efnb3*, ephrinB3 (rank #26/1192 in males; up-regulated; q=0.018; p=3.41E-05). *Sbf1*, SET Binding Factor 1 (rank #82/1192 in males; down-regulated; q=0.067; p=0.00039). *Fabp7*, Fatty Acid Binding Protein 7 (rank #84/1192 in males; up-regulated; q= 0.0725; p=0.00043). **d.** *Tcf7l2*, transcription factor 7 like 2 (rank #795/1192; up-regulated; q= 0.514; p=0.029). **e.** *Nf1*, Neurofibromin 1 (rank #1189/1192 in males; down-regulated; q=0.59; p=0.0499). **f.** *PLP1*, Proteolipid Protein 1 (rank #64/389 in females; down-regulated; q=1; p=0.0054). Normalized qRT-PCR data (to *pgk1*) is presented as mean fold changes ± SEM, n=10 samples/group. *p<0.05, **p<0.01, ****p<0.001 (unpaired Student’s t test with Welch’s correction). **g-g’.** Cerebellum protein contents determined by Western blot in males, and normalized to GAPDH levels (n=7/group). From RNAseq: PSAP, prosaposin (rank #950/1192 in males; up-regulated; q=0.5329; p=0.0359). Cfl1, cofilin-1 (rank #27/1192 in males; up-regulated; q=0.0192; p=4.03E-05). hnRNPκ, heterogeneous nuclear ribonucleoprotein κ (rank #787; up-regulated; q=0.509; p=0.0284). *p<0.05, **p<0.01; ***p<0.005 (unpaired Student’s t test with Welch’s correction).

**Extended Data Figure 4.**
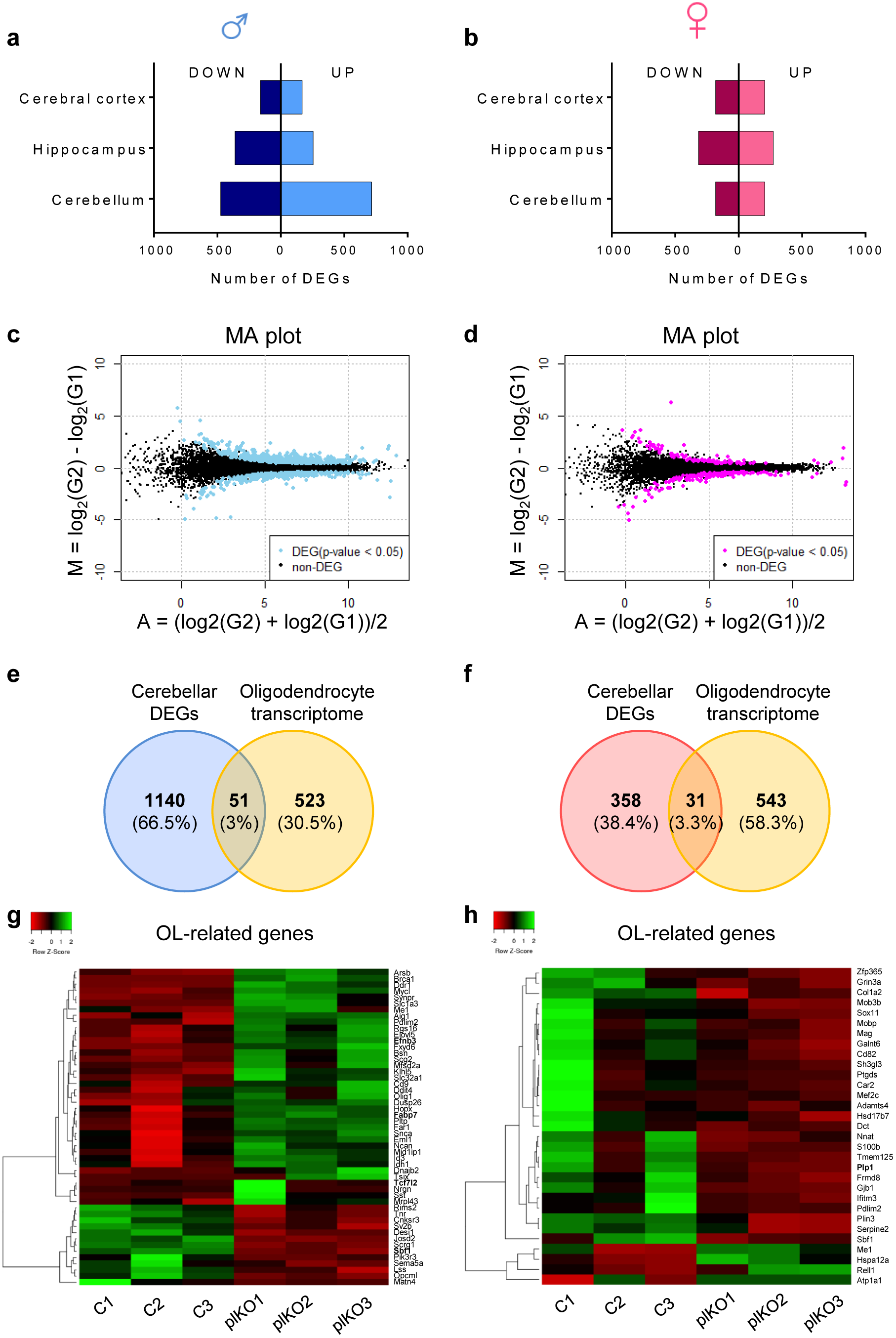
Long-term impact of placental ALLO insufficiency on brain transcriptome; highlight on the oligodendrocyte-associated genes. **a-b.** Bar graphs showing the number of differentially expressed genes (DEGs) that are up- or down-regulated in the cerebral cortex, hippocampus and cerebellum of plKO vs C mice at P30 in males (blue) and females (pink). The majority of DEGs were found in the male cerebellum. **c-d.** MA plots for the cerebellar RNAseq analysis of plKO vs C mice at P30. M, log ratio; A, mean average. DEGs are represented as colored dots (blue for males and pink for females). **e-f.** Venn diagrams showing the overlap between cerebellar DEGs in the plKO mice and the oligodendrocyte (OL) transcriptome ^13^. **g-h.** Heatmaps of OL-related cerebellar DEGs in C and plKO samples. Genes and samples were hierarchically clustered based on Pearson distance of z-score data and average linkage.

**Extended Data Figure 5.**
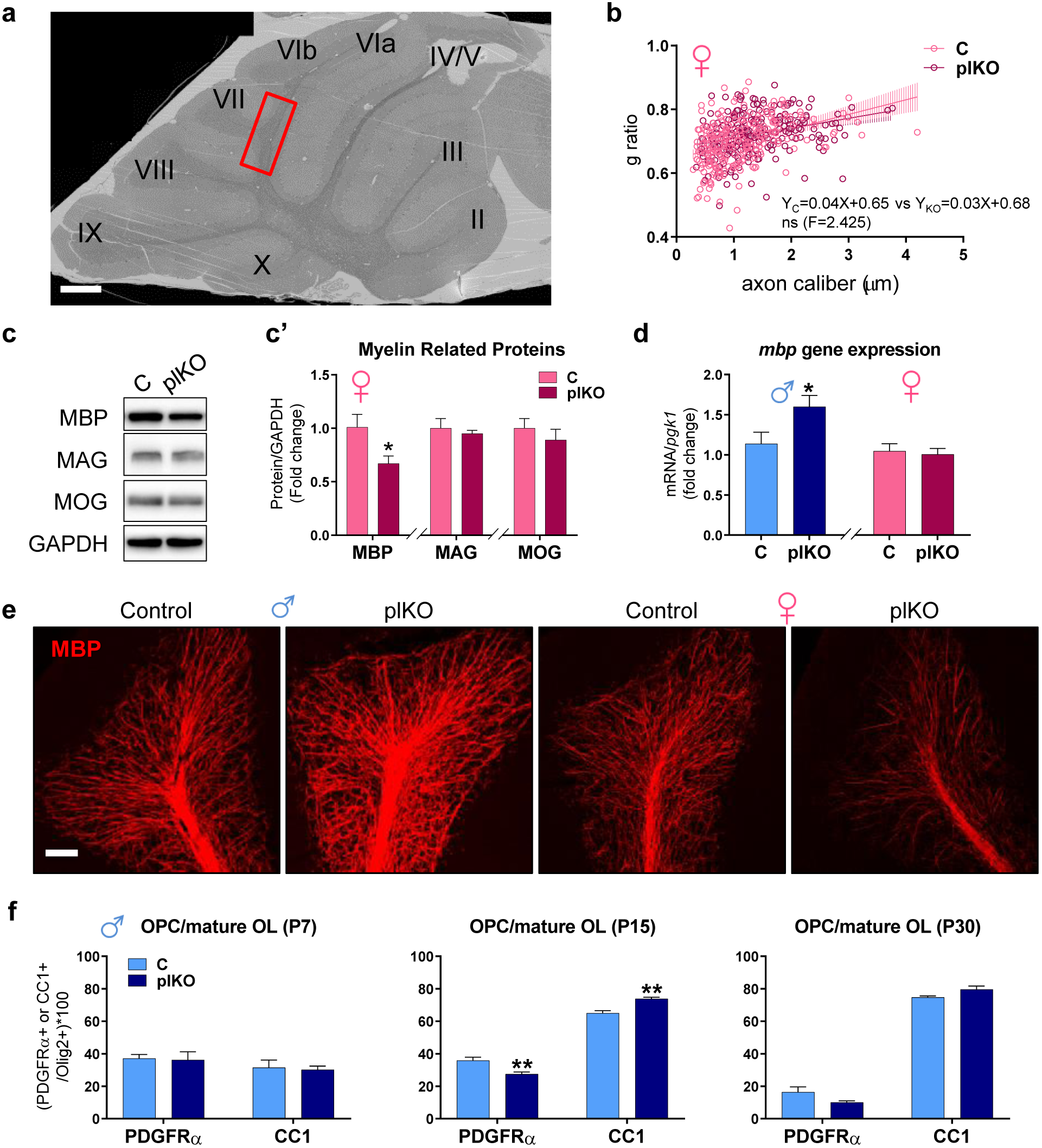
Minimal impact of placental ALLO insufficiency on cerebellar WM in females at P30. **a.** Scanning electron microscopy acquisition of a whole cerebellum ultrathin section showing the different cerebellar lobules (I to X) in a control mouse (650× low magnification). The region of interest (inter-lobule VI-VII) for high magnification acquisitions and g-ratio quantifications is indicated by red rectangle. Scale bar, 300 μm. **b.** Scatter plot of g ratios of >300 individual axons vs axon diameter; data from one representative C and one representative plKO female (light pink and dark pink, respectively). Fitted lines are linear regressions, showing unchanged g ratios in plKO vs C females. **c-c’.** Western blot analysis of myelin related proteins in the cerebellum of C and plKO mice. Cerebellar MBP contents are significantly lower in plKO than in C females. Normalized data to GAPDH is presented as means ± SEM (n=6/group). *p<0.05 (two-tailed unpaired Student’s t test with Welch’s correction). **d.** *mbp*-mRNA levels in the cerebellum of C and plKO mice. A significant up-regulation of MBP gene expression is evidenced in the plKO males only. Normalized qRT-PCR data is presented as mean fold changes ± SEM (n=11 samples/group). *p<0.05 (two-way ANOVA with Sidak’s multiple comparisons test). **e.** Immunofluorescent staining of MBP in the cerebellar lobule VII of C and plKO at P30. Sagittal 40-μm-thick section through the vermis. Laser scanning confocal microscopy (Leica TCS SP8). Scale bar, 100 μm. **f.** Ratios of OPCs (PDGFRα+ cells) or mature OLs (CC1+ cells) within the total oligodendrocyte lineage (Olig2+ cells) in the cerebellar WM across postnatal period. At P15, a transient acceleration of the OL maturation is observed in the male plKO, when cerebellar myelination is initiated (n=3-8/group). Data is presented as means ± SEM. **p<0.005 (two-way ANOVA with Sidak’s multiple comparisons test).

**Extended Data Figure 6.**
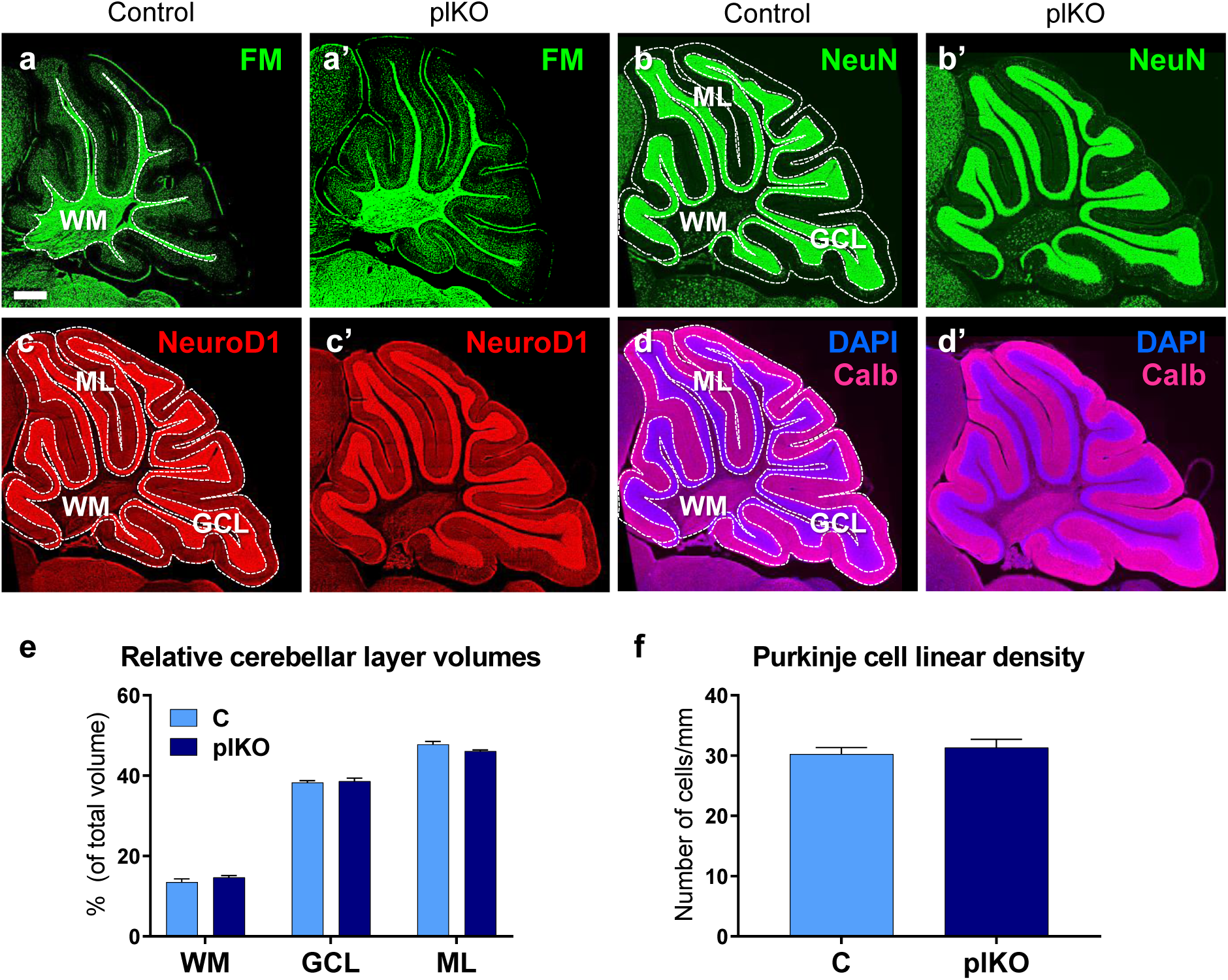
Histology of cerebellar grey and white matters in C and plKO males at P30. Cerebellar sagittal 40-μm-thick sections through the vermis. **a-a’.** FluoroMyelin (FM) green staining. **b-b’.** NeuN-immunofluorescence (marker for neuronal nuclei). **c-c’.** NeuroD1-immunofluorescence (marker for granule cells). **d-d’.** Calbindin-immunofluorescence (Purkinje cell marker). **e.** Cerebellar layer volumes were unchanged in plKO when compared with C mice. Data is presented as means ± SEM from 4 sections/animal and 6-7 animals/genotype (two-way ANOVA with Sidak’s multiple comparisons test). **f.** Purkinje cell linear density in the lobule VI-VII of C and plKO males at P30. Data is presented as means ± SEM from 4 sections/animal and 6-7 animals/genotype (p=0.7308; (two-tailed unpaired Student’s t test with Welch’s correction). Calb, calbindin; GCL, granule cell layer; ML, molecular layer; WM, white matter.

**Extended Data Figure 7.**
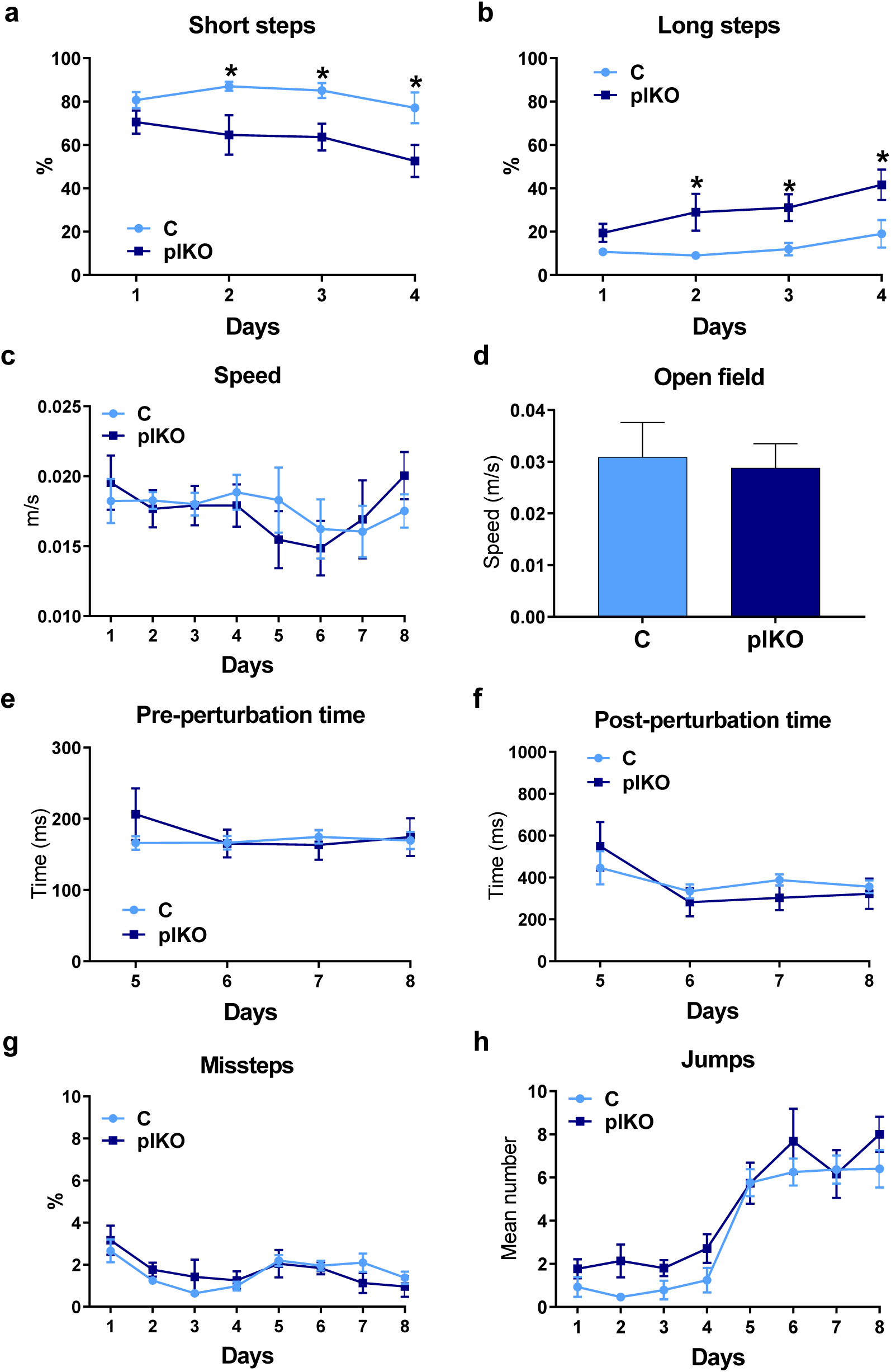
Performance of male offspring on the Erasmus ladder test at P30. **a-b.** The analysis of baseline gait and coordination on days 1-4 showed significantly increased step length in adult male plKO mice than in their control littermates. Data is presented as means ± SEM (n=8 C and 6 plKO). *p<0.05 (two-way repeated-measures ANOVA with Sidak’s multiple comparisons test). **c.** Speed on the ladder was unchanged between C and plKO mice (two-way repeated-measures ANOVA with Sidak’s multiple comparisons test). **d.** Locomotor speed was unchanged in the open-field (n=6-8/group; two-tailed unpaired Student’s t test with Welch’s correction). **e-h.** The associative cerebellar learning was tested with the introduction of a conditioned tone and obstacle on days 5-8. The plKO mice exhibited no motor learning deficit. Data is presented as means ± SEM, n=8 C and 6 plKO, *p<0.05 (two-way repeated-measures ANOVA with Sidak’s multiple comparisons test).

**Extended Data Figure 8.**
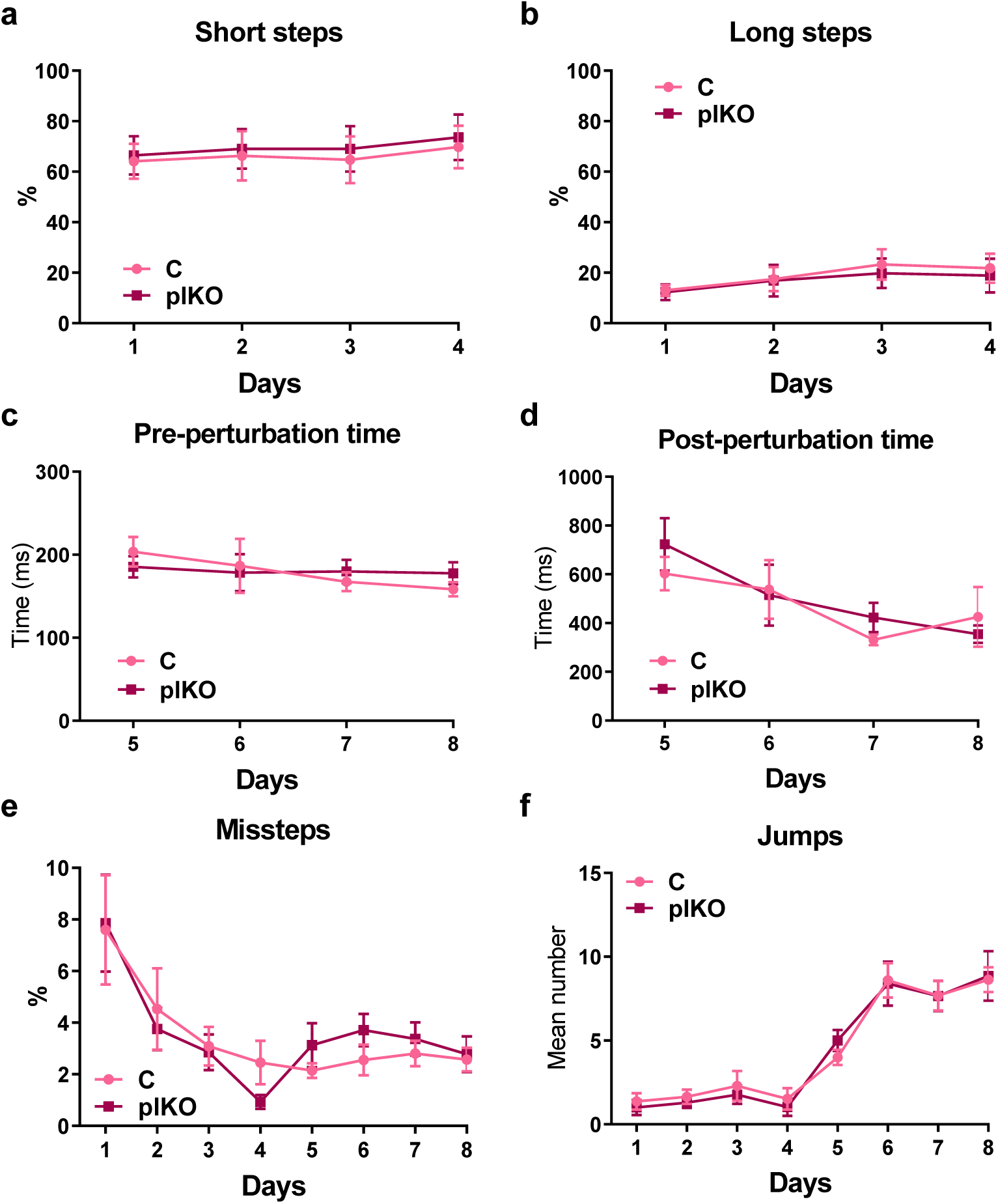
Performance of female offspring on the Erasmus ladder test at P30. **a-b.** The analysis of baseline gait and coordination on days 1-4 showed no difference in female plKO mice vs their control littermates (two-way repeated-measures ANOVA with Sidak’s multiple comparisons test). **c-f.** The associative cerebellar learning was tested with the introduction of a conditioned tone and obstacle on days 5-8. The plKO mice exhibited no motor learning deficit. Data is presented as means ± SEM, n=7 C and 8 plKO (two-way repeated-measures ANOVA with Sidak’s multiple comparisons test).

**Extended Data Figure 9.**
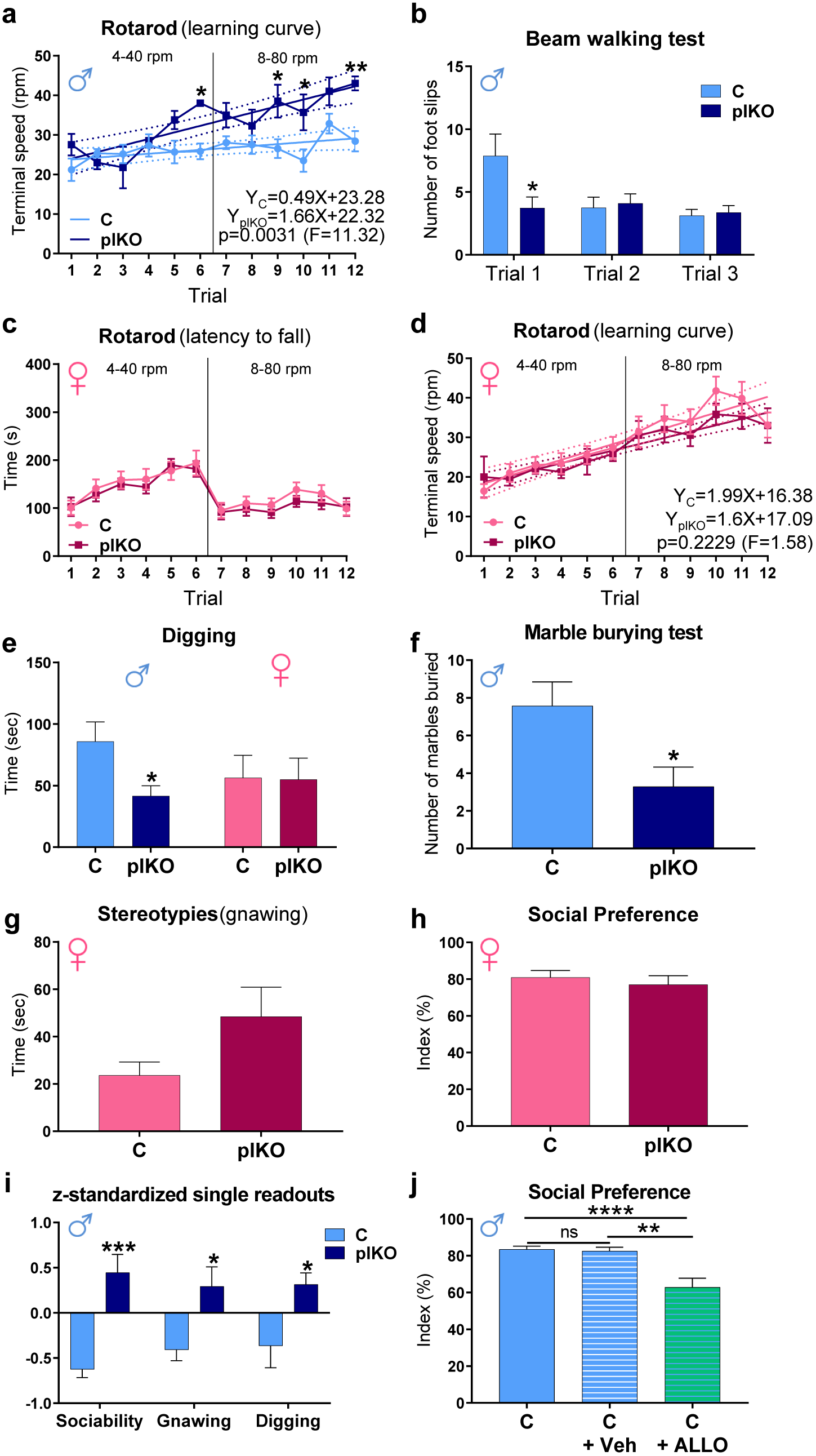
Sex-dependent behavioural alterations associated with placental ALLO insufficiency at P30. **a.** Performance of male C vs plKO littermates on the accelerating rotarod. Terminal speed of rotation, presented at 4 to 40 rpm (trials 1-6) and 8 to 80 rpm (trials 7-12), was used to calculate learning rate (linear regression). plKO males exhibit higher learning rate than their C littermates (n=6 C and 4 plKO). Data is presented as means ± SEM; *p<0.05, **p<0.01; significant difference between groups (two-way repeated-measures ANOVA with Sidak’s multiple comparisons test). **b.** Performance on the inclined beam walking test. plKO males make significant less foot slips than the controls during the first trial (n=16 C and 11 plKO). Data is presented as means ± SEM; *p<0.05; significant difference between groups (two-way repeated-measures ANOVA with Sidak’s multiple comparisons test). **c.** Performance of female C vs plKO littermates on the accelerating rotarod. Time to fall off is presented at 4 to 40 rpm and 8 to 80 rpm. Data is presented as means ± SEM, n=9 C and 7 plKO (two-way repeated-measures ANOVA with Sidak’s multiple comparisons test). **d.** Terminal speed of rotation on the accelerating rotarod, presented at 4 to 40 rpm (trials 1-6) and 8 to 80 rpm (trials 7-12), was used to calculate learning rate (linear regression). No difference was evidenced between C and plKO female littermates. Data is presented as means ± SEM, n=9 C and 7 plKO (two-way repeated-measures ANOVA with Sidak’s multiple comparisons test). **e.** Time of spontaneous digging over a 15-min period in males (blue) and females (pink). Male plKO mice spend less time digging than their C littermates (Males: n=20 C and 28plKO; Females: n=10 C and 10 plKO). Data is presented as means ± SEM; *p<0.05; significant difference between groups (two-way ANOVA with Sidak’s multiple comparisons test). **f.** Marble burying test. Graph represents the number of marbles at least 2/3^rd^ buried over 30-min (n=7/group). Data is presented as means ± SEM; *p<0.05; significant difference between groups (Mann-Whitney test). **g.** Spontaneous repetitive behaviour (gnawing) observed over a 15-min period of time in females. Data is presented as means ± SEM (n=10 C and 9 plKO (two-tailed unpaired Student’s t test with Welch’s correction). **h.** 3-chamber sociability test in females. Data is presented as means ± SEM (n=9 C and 6 plKO; two-tailed unpaired Student’s t test with Welch’s correction). **i.** z-standardized single significant behaviour readouts that are integrated into the autism composite score (n=20-28/group). Data is presented as means ± SEM; *p<0.05; ***p<0.0005; significant difference between groups (two-way ANOVA with Sidak’s multiple comparisons test). **j.** 3-chamber sociability test. ALLO injection to the dams at E15.5 significantly alters social preference in the male C mice (n=20 C, 9 C+Veh and 14 C+ALLO). Social Preference Index was calculated as above. Data is presented as means ± SEM; **p<0.005; ****p<0.0001 significant difference between groups (one-way ANOVA with Tukey’s multiple comparisons test).

